# Enhancing adult neuroplasticity by epigenetic regulation of Parvalbumin-expressing GABAergic cells

**DOI:** 10.1101/2022.08.09.503382

**Authors:** Marisol Lavertu-Jolin, Bidisha Chattopadhyaya, Pegah Chehrazi, Denise Carrier, Florian Wünnemann, Séverine Leclerc, Félix Dumouchel, Derek Robertson, Hicham Affia, Kamal Saba, Vijaya Gopal, Anant Bahadur Patel, Gregor Andelfinger, Graçiela Pineyro, Graziella Di Cristo

**Affiliations:** Centre de Recherche, CHU Sainte-Justine (CHUSJ), Montréal, Canada; Department of Neurosciences, Université de Montréal, Montréal, Canada; Department of Pharmacology, Université de Montréal, Montréal, Canada; Department of Pediatrics, Université de Montréal, Montréal, Canada; CSIR-Centre for Cellular and Molecular Biology, Hyderabad 500007, India; Academy of Scientific and Innovative Research, Ghaziabad 201002, India; Heidelberg University, Faculty of Medicine & Heidelberg University Hospital, Institute for Computational Biomedicine, Bioquant, Heidelberg, Germany

## Abstract

Failure of inhibiting fear in response to harmless stimuli contributes to anxiety disorders. Extinction training only temporarily suppresses fear memories in adults, but it is highly effective in juveniles. GABAergic parvalbumin-positive (PV^+^) cells restrict plasticity in adult brains, thus increasing PV^+^ cell plasticity could promote the suppression of fear memories following extinction training in adults. Histone deacetylase 2 (Hdac2) restrains both structural and functional synaptic plasticity; however, whether and how Hdac2 controls adult PV^+^ cell plasticity is unknown. Here, we report that *Hdac2* deletion or pharmacological inhibition in PV^+^ cells attenuate spontaneous recovery of fear memory after fear extinction learning in adults. These manipulations promote a temporally restricted downregulation of *Acan*, a critical perineuronal net component expressed exclusively by PV^+^ cells in medial prefrontal cortex. Finally, we show that *Acan* transient downregulation before extinction training but after fear memory acquisition is sufficient to reduce spontaneous fear memory recovery in wild-type mice.

## Introduction

Anxiety and trauma-related disorders are associated with a huge socioeconomic burden, given the limited treatment options available^1^. Certain anxiety disorders are characterized by abnormally persistent emotional memories of fear-related stimuli, and impaired inhibition of learned fear is a feature in post-traumatic stress disorder (PTSD), anxiety disorders and phobias^2^. In rodents, pairing a neutral tone (conditioned stimulus, CS) with an aversive foot shock (unconditioned stimulus, US) leads to the formation of a robust fear memory that can last an entire lifetime^3^. The reduction of fear responses through fear extinction learning is at the heart of clinical exposure psychotherapies applied in the case of PTSD^4,5^. In adults, extinction learning, or the gradual decrease of behavioural response to a CS that occurs when the stimulus is presented without reinforcement (e.g., aversive foot shock), is neither robust nor permanent since conditioned fear responses can recover spontaneously with the passage of time; in other words, extinction learning is unstable in adults^3,6,7^. Strategies aimed at enhancing the ability to inhibit responses to associations that are no longer relevant may have strong therapeutic value. Several studies suggested that the maturation of GABAergic neurotransmission contributes to the reduced long-term retention of extinction learning in adults^8,9^. Therefore, modulating GABAergic function is an attractive target to reduce spontaneous recovery of fear memories with time after extinction training in adults. However, global or prolonged reduction of GABAergic drive can generate undesirable effects such as epileptic activity. Therefore, ideal approaches targeting modulation of GABAergic transmission to foster adult brain plasticity need to be both cell type-specific and temporally-controlled.

Amongst the different GABAergic interneuron types, reducing parvalbumin-expressing (PV^+^) interneuron activity or connectivity is sufficient to reinstate heightened plasticity in adult sensory cortices^10-13^. Recent studies showed that PV^+^ cells also play a role in fear memory extinction. First, fear responses during extinction learning are increased by chemogenetic activation of PV^+^ cells in the medial prefrontal cortex^14^. Second, extinction training induces remodelling of perisomatic PV^+^ synapses around excitatory neurons that were previously activated during fear conditioning in the basolateral amygdala (BLA)^15^. Third, PV^+^ cell-specific deletion of Nogo Receptor 1, a neuronal receptor for myelin-associated growth inhibitors^12^, enhances both BLA PV^+^ synapse remodeling and the retention of fear extinction^16^. Fourth, fear extinction weakens the functional outputs of PV^+^ cells to pyramidal neurons in medial prefrontal cortex^14^. All together, these data suggest GABAergic PV^+^ cell plasticity, an activity-regulated process^17-19^, is required for extinction learning and retention.

Transcriptional mechanisms coupling synaptic activity to changes in gene expression drive cellular processes that mediate behavioural adaptations^20^. Accessibility of the chromatin to activate these transcriptional programs is modulated by changes in the state of histone acetylation of chromatin. In particular, blocking endogenous histone deacetylases (Hdacs) promotes both structural and functional synaptic plasticity processes^21,22^ that are required to modify behaviour and improve performance during learning. Accordingly, administration of pan-Hdac inhibitors in adult mice reactivates visual cortical plasticity^23^ and enhances extinction learning^24^. Several studies have shown that, of the many Hdacs expressed in the mammalian brain, Hdac2 plays a critical role in fear learning and memory^21,22,25^. Therefore, in principle it is conceivable that Hdac2 expression in PV^+^ cells constrain their plasticity, thus reducing the long-term retention of extinction learning in adults.

PV^+^ interneurons are the only cortical cell type surrounded by perineuronal nets (PNNs), which are lattice-like aggregates of chondroitin sulfate proteoglycans-containing extracellular matrix^19^. The percentage of PV^+^ interneurons enwrapped by PNN increases during development in different brain regions, including those involved in fear memory formation and extinction^19^. Recent evidences shows that PNNs stabilize afferent synaptic contacts onto mature cortical PV^+^ cells, thus limiting their plasticity^19,26,27^. Indeed, degradation of PNNs in adult BLA promotes fear memory erasure^28^, a phenomena observed only in younger animals^8,28,29^. PNNs are dynamic structures, comprising of numerous structural components and remodeled by metalloproteases^30^. Whether the expression of distinct PNN components is controlled by epigenetic regulation specifically in PV^+^ cells, in turn modulating PV^+^ cell plasticity, is unknown.

Here, we report that *Hdac2* deletion or pharmacological inhibition in PV^+^ cells attenuate spontaneous recovery of fear memory after fear extinction learning in adults. In particular, these manipulations promote a temporally restricted downregulation of *Acan*, coding for aggrecan, a critical perineuronal net component, which we show to be expressed almost exclusively by PV^+^ cells in medial prefrontal cortex. Finally, we show that intravenous siRNA against *Acan* delivery right before extinction induces a brief knock-down of *Acan* expression and reduces spontaneous fear recovery with time in adult wild-type mice. Therefore, the transient downregulation of *Acan* expression before extinction training but after fear memory acquisition is sufficient to reduce spontaneous fear memory recovery in wild-type mice.

## Results

### Hdac2 inhibition, in PV^+^ cells, leads to reduced spontaneous recovery of fear memory with time following extinction training in adult mice

To evaluate whether and how *Hdac2* deletion specifically in PV^+^ cells affects fear memory formation, extinction and spontaneous recovery, we crossed mice carrying a conditional allele of *Hdac2* (*Hdac2*^*lox/lox* 21^), which allows cell-specific developmental stage-restricted manipulation of *Hdac2*, with mice expressing Cre recombinase under the control of the PV promoter (*PV-Cre*, ^31^). In cortical GABAergic cells, PV starts to be expressed after the first postnatal week and peaks after the third week. This breeding scheme generated PV^+^-cell restricted homozygous (*PV-Cre*^*+/−*^;*Hdac2*^*lox/lox*^) mice and control *PV-Cre*^*−/−*^ littermates (*PV-Cre*^*−/−*^;*Hdac2*^*lox/lox*^ and *PV-Cre*^*−/−*^;*Hdac2*^*lox/+*^ mice, referred to hereafter as *Hdac2*^*lox*^). To confirm the specificity of Cre expression in *PV-Cre*^*+/−*^ *;Hdac2*^*lox/lox*^ mice, we bred them with *RCE*^*GFP*^ reporter mice^32^ and quantified GFP^+^/PV^+^ colocalized cells in somatosensory cortex (SSCX), medial prefrontal cortex (PFC) and basolateral amygdala (BLA). At P60, we observed that all GFP^+^ cells in all brain regions studied expressed PV, whereas the recombination rate was region specific (GFP^+^PV^+^/PV^+^ cells: SSCX: 82 ± 5 %, n=3 mice; PFC: 58 ± 8 %, n=5 mice; BLA: 38 ± 3 %, n=3 mice). Co-immunolabelling of PV and Hdac2 in coronal brain sections confirmed that Hdac2 expression levels were significantly reduced in PV^+^ cells in the conditional KO mice compared to their control littermates (Supplementary Fig. 1). We then assessed fear memory formation, extinction rate and spontaneous recovery in *PV-Cre*; *Hdac2*^*lox/lox*^ mice (Fig. 1a). After fear conditioning, both *PV-Cre*; *Hdac2*^*lox/lox*^ and *Hdac2*^*lox*^ mice presented strong freezing behaviour, suggesting that fear learning was not affected by *Hdac2* deletion in PV^+^ cells (freezing time at early extinction CS1-2, Mann-Whitney test, *P* = 0.1878, Fig. 1b). Compared to wild-type littermates, conditional KO mice showed similar extinction rate during training, but significantly less freezing during the retrieval and renewal tests 7 days later (Fig. 1b), suggesting reduced spontaneous recovery of fear memory. The reduced fear expression was not due to loss of memory, since in absence of extinction training, freezing levels were undistinguishable between conditional KO and control mice 10 days after fear conditioning (Fig. 1c). Further, this was not due to altered motor or anxiety behavior, since we found no differences between the genotypes in the open field and elevated plus maze assays (Fig. 1d, e). To ensure that the observed phenotype was caused by the genetic deletion of *Hdac2* rather than any side-effect of the *Cre* insertion in the *PV* locus^31^, we generated a second mouse line in which control littermates also carried the *PV-Cre* allele, while the *Hdac2* locus remained wild-type (*PV-Cre*^*+/−*^*;Hdac2*^*+/+*^ mice). Consistent with our previous observations, these conditional KO mice showed significantly reduced freezing during the retrieval test compared to their control littermates (Supplementary Fig. 2a). In absence of extinction training, both genotypes showed strong fear memory 10 days after fear learning (Supplementary Fig. 2b). Overall, these data suggest that the deletion of *Hdac2* restricted to postnatal PV^+^ cells is sufficient to improve fear extinction retention over time in adults.

**Figure 1.**
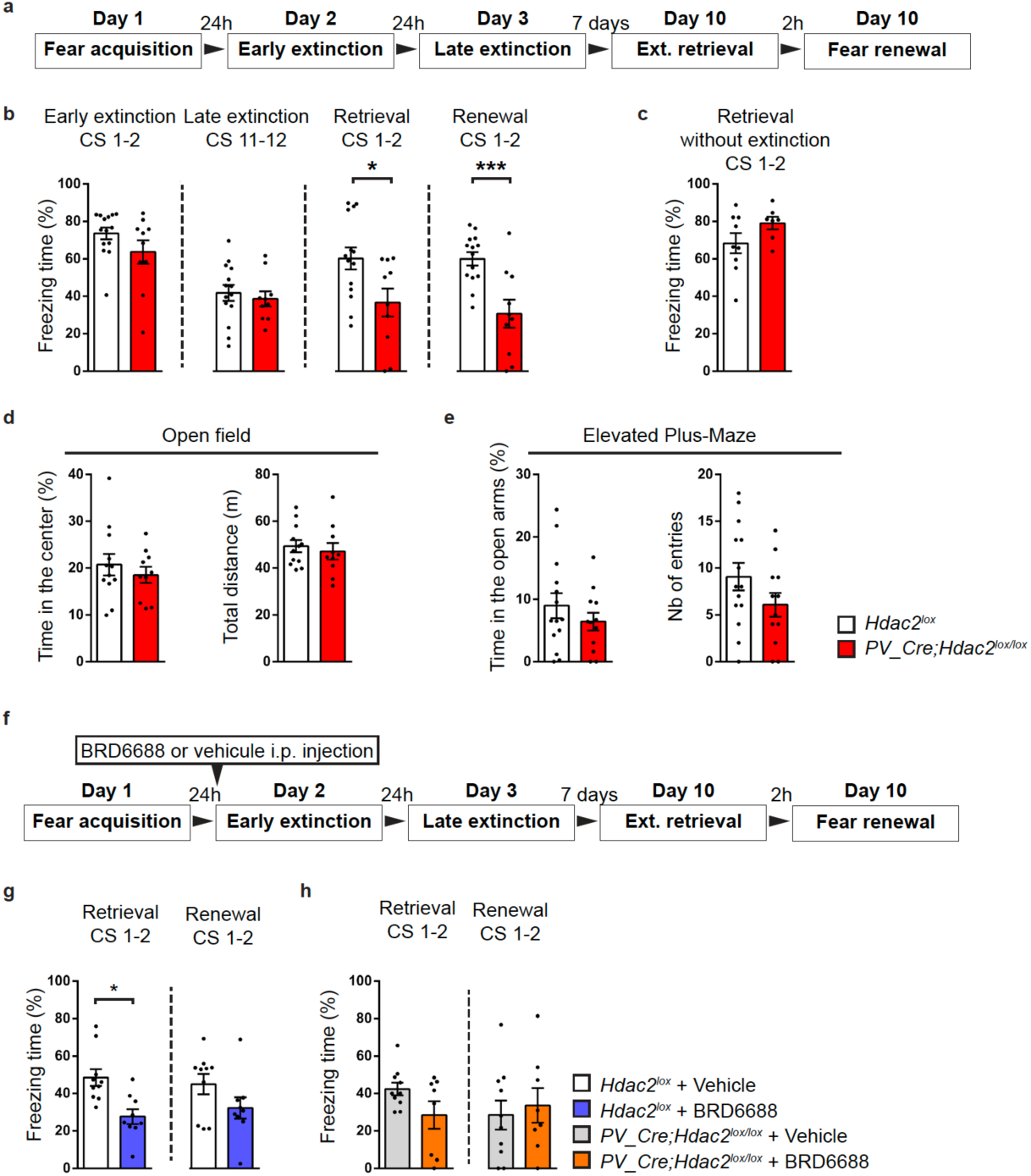
Postnatal *Hdac2* deletion restricted to PV^+^ cells or Hdac2 inhibition before extinction training decrease the spontaneous recovery of fear memory after extinction in adult mice. (**a**) Schematic representation of the experimental protocol. Ext.: extinction. (**b**) Wild-type and *PV-Cre;Hdac2*^*lox/lox*^ littermates show efficient fear extinction. Repeated two-way ANOVA; F_genotype_ (1,22)=4.430, *P*=0.0470, F_extinction_(11,242)=8.864, *P*<0.0001, F_genotype_*_extinction_ (11,242)=0.6427, *P*=0.7911. Wild-type, but not *PV-Cre;Hdac2*^*lox/lox*^ littermates, show spontaneous recovery and context-dependent renewal of fear memory. Unpaired two-tailed *t*-tests, *P*=0.0205 for fear retrieval; *P*=0.0008 for fear renewal. Number of mice: *Hdac2*^*lox/+*^ or *Hdac2*^*lox/lox*^ n=14; *PV-Cre;Hdac2*^*lox/lox*^ n=10. (**c**) In the absence of extinction training, *PV-Cre;Hdac2*^*lox/lox*^ mice show stable fear memory 10 days after conditioning, as control littermates (Mann-Whitney test, *P*=0.2509). Number of mice: *Hdac2*^*lox/+*^ or *Hdac2*^*lox/lox*^ n=9; *PV-Cre;Hdac2*^*lox/lox*^ n=7. (**d**) In the open field test, there is no significant difference in time spent in the center of the open field (unpaired two-tailed *t*-test, *P*=0.4635) nor the total distance travelled (unpaired two-tailed *t*-test, *P*=0.6083) between the two genotypes. Number of mice: *Hdac2*^*lox/+*^ or *Hdac2*^*lox/lox*^ n=12; *PV-Cre;Hdac2*^*lox/lox*^ n=10. (**e**) In the elevated plus maze test, there is no genotype-dependent difference in time spent (unpaired two-tailed *t*-test, *P*=0.3279) nor the number of entries in the open arms (unpaired two-tailed *t*-test, *P*=0.1449). Number of mice: *Hdac2*^*lox/+*^ or *Hdac2*^*lox/lox*^ n=14; *PV-Cre;Hdac2*^*lox/lox*^ n=12. (**f**) Schematic representation of the experimental protocol. Hdac2 inhibitor BRD6688 or the vehicle was injected intraperitoneally (i.p.) 6 hours before the early extinction procedure. (**g**) One week after extinction training, freezing levels are significantly different in the spontaneous recovery test between vehicle- and BRD6688-injected *Hdac2*^*lox/lox*^ (**h**) but not between vehicle- and BRD6688-injected *PV-Cre;Hdac2*^*lox/lox*^ mice. (**g**) Extinction retrieval unpaired two-tailed *t*-test, *P*=0.0030; fear renewal, unpaired two-tailed *t*-test, *P*=0.1246. Number of *Hdac2*^*lox/lox*^ mice injected with vehicle, n=10 or BRD6688, n=9. (**h**) Extinction retrieval unpaired two-tailed *t*-test *P*=0.0850, fear renewal unpaired two-tailed *t*-test *P*=0.6733. Number of *PV-Cre;Hdac2*^*lox/lox*^ mice injected with vehicle, n=10 or BRD6688, n=8. Graph bars represent mean ± s.e.m. Circles represent individual mouse values. * *P* < 0.05, *** *P* < 0.001.

Reduced spontaneous recovery of fear memories over time, after extinction training, in adult *PV-Cre*;*Hdac2*^*lox/lox*^ mice could rely on different mechanisms recruited during fear memory acquisition in the conditional KO mice compared to control littermates, as suggested by the increased fear memory erasure susceptibility only when the enzymatic degradation of PNN in the BLA of adult mice is performed before fear memory acquisition^28^. Alternatively, *Hdac2* deletion in PV^+^ cells may specifically affect the retention of extinction memories. To distinguish between these two hypotheses, we used BRD6688, a small molecule with kinetic selectivity for Hdac2 as compared to Hdac1, a close family member showing 92% homology within the catalytic domain with Hdac2^33,34^. We first asked whether pharmacological inhibition of Hdac2 after fear learning but before the onset of extinction training was sufficient to reduce spontaneous fear memory recovery (Fig. 1f). Compared to vehicle-injected mice, BRD6688-injected mice presented less freezing behavior at the retrieval test (Fig. 1g). Therefore, combining a brief Hdac2 inhibition with extinction training rendered the extinction memory stronger, increasing its persistence. Circuits other than PV^+^ cells may contribute to the promoting effect of Hdac2 inhibition on the long-term retention of extinction memories. However, BRD6688 treatment could not further reduce freezing behavior in *PV-Cre; Hdac2*^*lox/lox*^ mice during retention behaviour (Fig. 1h), thus supporting the hypothesis that the state of chromatin acetylation regulated by Hdac2 in PV^+^ cells plays a critical role in limiting the spontaneous recovery of fear memory with time after extinction training.

### Postnatal deletion of *Hdac2* in PV^+^ cells enhances the remodelling potential of their synaptic terminals

During their maturation process, PV^+^ cells form numerous perisomatic synapses on pyramidal cell somata over time^35^. To investigate whether *Hdac2* conditional deletion in PV^+^ cells affected their efferent connectivity, we quantified putative perisomatic synapses formed by PV^+^ cells onto pyramidal cells by co-immunolabeling with gephyrin (scaffolding protein associated with GABA-A receptor, postsynaptic) and PV, in PFC and BLA. We found that the density of PV^+^ boutons colocalizing with gephyrin was significantly reduced in both PFC and BLA of naive conditional KO mice compared to their control littermates (Fig. 2a, b). This difference was not due to decreased PV expression, since the density of perisomatic PV^+^ boutons was not significantly different between the two genotypes (Fig. 2b).

**Figure 2.**
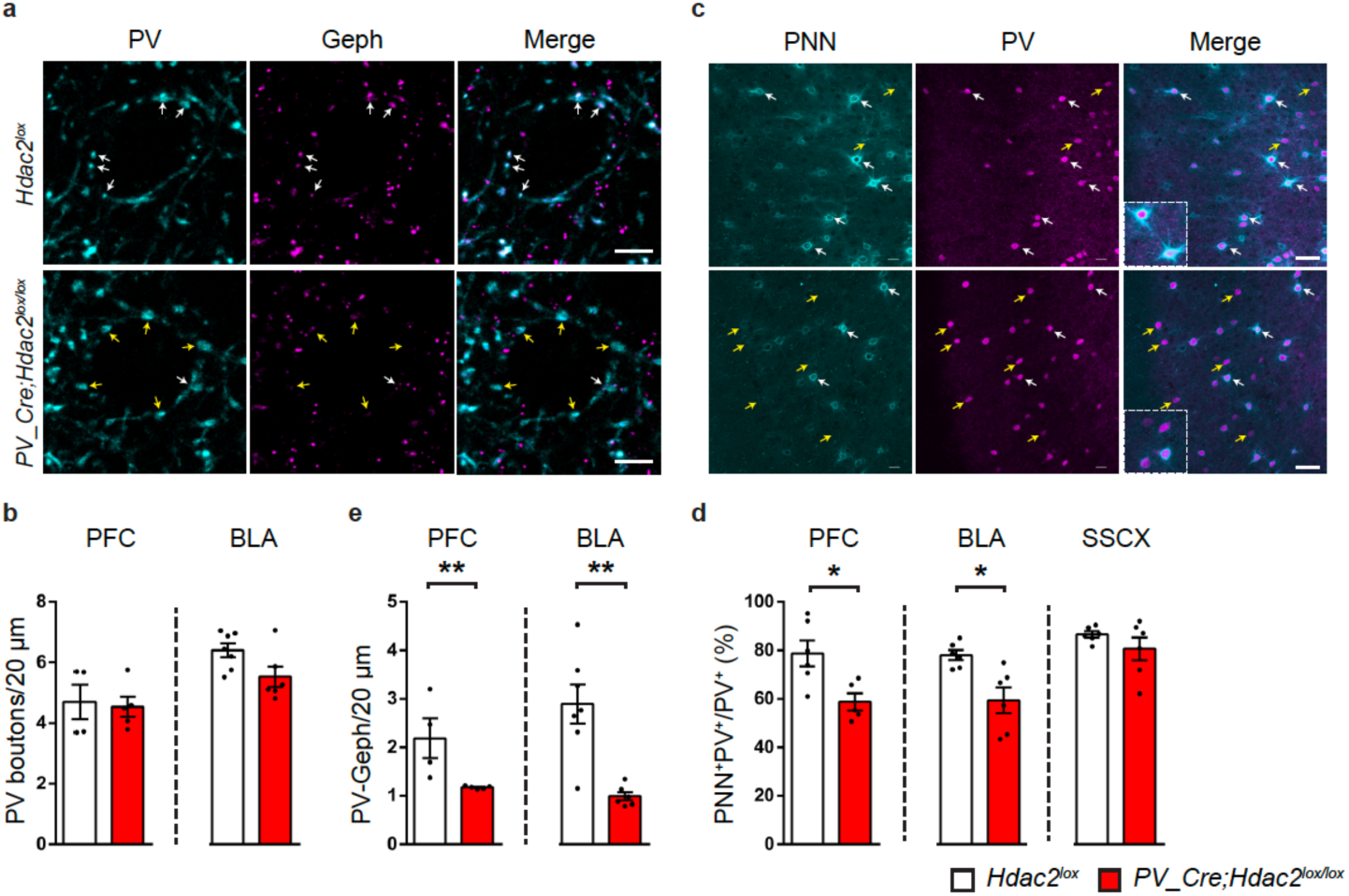
*PV_Cre;Hdac2*^*lox/lox*^ mice show decreased PV^+^ cell efferent synaptic connectivity and reduced PNN agglomeration around cortical PV^+^ cell bodies. (a) PFC coronal sections of *PV-Cre;Hdac2*^*lox/lox*^ and control littermates immunolabeled for PV (cyan) and gephyrin (magenta). White arrows indicate perisomatic PV^+^Gephyrin^+^ boutons, yellow arrows indicate PV^+^Gephyrin^-^ boutons. Scale bar, 5 µm. (**b**) The density of perisomatic PV^+^ boutons is not significantly different neither in PFC nor in BLA between the two genotypes. PFC: Mann-Whitney test, *P*=0.6825, *Hdac2*^*lox/lox*^ n=4; *PV-Cre;Hdac2*^*lox/lox*^ n=5;. BLA: Mann-Whitney test, *P*=0.1014, *Hdac2*^*lox/lox*^ n=7; *PV-Cre;Hdac2*^*lox/lox*^ n=6. (**c**) The density of perisomatic PV^+^ boutons co-localizing with gephyrin puncta is significantly reduced in both PFC and BLA of conditional KO mice compared to their littermates. PFC: Mann-Whitney test, *P*=0.0159, *Hdac2*^*lox/lox*^ n=4; *PV-Cre;Hdac2*^*lox/lox*^ n=5. BLA: Mann-Whitney test, *P*=0.0140, *Hdac2*^*lox/lox*^ n=7; *PV-Cre;Hdac2*^*lox/lox*^ n= 6). (**d**) Coronal sections of PFC of *PV-Cre;Hdac2*^*lox/lox*^ and control littermates labelled with WFA (PNN, cyan) and PV (magenta). White arrows indicate PV^+^PNN^+^ cell bodies while yellow arrows point to PV^+^PNN^-^ cell bodies. Scale bar, 40 µm. (**e**) The percentage of PV^+^ cell bodies surrounded by WFA-stained PNN is significantly reduced in the PFC and BLA, but not in the SSCX, of conditional KO mice compared to their littermates. PFC: Mann-Whitney test, *P*=0.0173, *Hdac2*^*lox/lox*^, n=6; *PV-Cre;Hdac2*^*lox/lox*^, n=5. BLA: Mann-Whitney test, *P*=0.0087, *Hdac2*^*lox/lox*^, n=6; *PV-Cre;Hdac2*^*lox/lox*^, n=6. SSCX: Mann-Whitney test, *P=*0.6753, *Hdac2*^*lox/lox*^, n=6; *PV-Cre;Hdac2*^*lox/lox*^ n=6. Data represent mean ± s.e.m. Black circle represent individual data points. * P < 0.05, ** *P* < 0.01.

Previous reports showed that remodeling of perisomatic inhibitory synapses was critical for efficient extinction of fear memories^15,16^. To investigate whether PV^+^ cell remodelling potential was affected by *Hdac2* deletion, we analyzed PV^+^ cell perisomatic synapses in both PFC and BLA 24 hrs after the end of fear extinction training in *PV-Cre; Hdac2*^*lox/lox*^ mice and their control littermates. While naive conditional mutant mice showed decreased percentage of perisomatic PV^+^ boutons colocalizing with gephyrin puncta compared to control mice in both PFC and BLA (Fig. 2c, Supplementary Fig. 3a, b, e), no significant difference could be detected between the two genotypes following extinction training (Supplementary Fig. 3c, d, e). These results suggest that fear extinction training promoted more extensive PV^+^ cell synapse remodelling in *PV-Cre; Hdac2*^*lox/lox*^ mice than in control littermates, reinforcing the hypothesis that changes in PV^+^ cell inhibitory drive is critical for the long-term retention of extinction memory^15^.

### Postnatal deletion of *Hdac2* in PV^+^ cells reduces PNN aggregation around their somata

PNN appearance around PV^+^ cell somata and dendrites is developmentally regulated, and correlates with increased inhibition and critical period closure in sensory cortices^36^. PV^+^ cell enwrapping by PNN has been shown to be a limiting factor for adult brain plasticity^19,26,27^. In particular, chondroitin sulfate proteoglycans assembly into PNNs promotes the formation of extinction-resistant memories^28^. Since we observed reduced spontaneous recovery of fear memory over time following extinction training in adult *PV-Cre*; *Hdac2*^*lox/lox*^ mice, we reasoned that this process may be accompanied by PNN reduction in conditional KO mice. Using WFA staining to label PNNs, we indeed found a significant decrease in the percentage of PV^+^ cells surrounded by PNNs in PFC and BLA, two regions directly implicated in fear behaviour regulation, in naive mutant mice compared to control littermates (Fig. 2c, d). Of note, we observed that PNNs were not localized exclusively around PV^+^ cells in the BLA (data not shown), consistent with previous reports^37,38^. Unexpectedly, we did not observe a significant difference in the percentage of PNN^+^PV^+^ cells in the SSCX of *PV-Cre*; *Hdac2*^*lox/lox*^ compared to *Hdac2*^*lox*^ mice (Fig. 2d), suggesting that epigenetic regulation of PNN organization is region-specific in the cortex.

### Hdac2 regulates Aggrecan expression in PV^+^ cells

WFA specifically binds to CAG-chains^39^ in all lecticans. Since we observed decreased WFA labeling surrounding PV^+^ cell somata in brain regions involved in fear regulation in *PV-Cre*; *Hdac2*^*lox/lox*^ mice (Fig. 2c, d), we next sought to identify which lecticans are cell-autonomously expressed by PV^+^ cells, and could thus be directly affected by PV^+^ cell-specific *Hdac2* deletion. Using Drop-Seq, we transcriptionally profiled PFC cells from 6 weeks-old wild-type mice. Samples from 5 biological replicates yielded 19393 cells that were sequenced to a median depth of 15,555 aligned reads/cell (representing a median of 1427 genes/cell). Cells were then clustered and represented in t-Distributed Stochastic Neighbor Embedding (t-SNE) plots (Fig. 3a). Cell clusters were defined as glutamatergic neurons, GABAergic neurons or non-neuronal cells based on *Slc17a7, Gad1* and *Gad2* detection (Supplementary Fig. 4). Using the Dropviz database as a template for murine cortical transcriptomes, clusters were then associated with well-known neuronal and non-neuronal cell groups. Among the latter, we distinguished astrocytes, oligodendrocytes/polydendrocytes, microglial cells/macrophages as well as endothelial cells. Within neuronal cell types, VGLUT^+^ glutamatergic neurons and Gad1^+^/Gad2^+^ interneurons were identified, including PV-expressing interneurons (cluster 9, Supplementary Fig.5).

**Figure 3.**
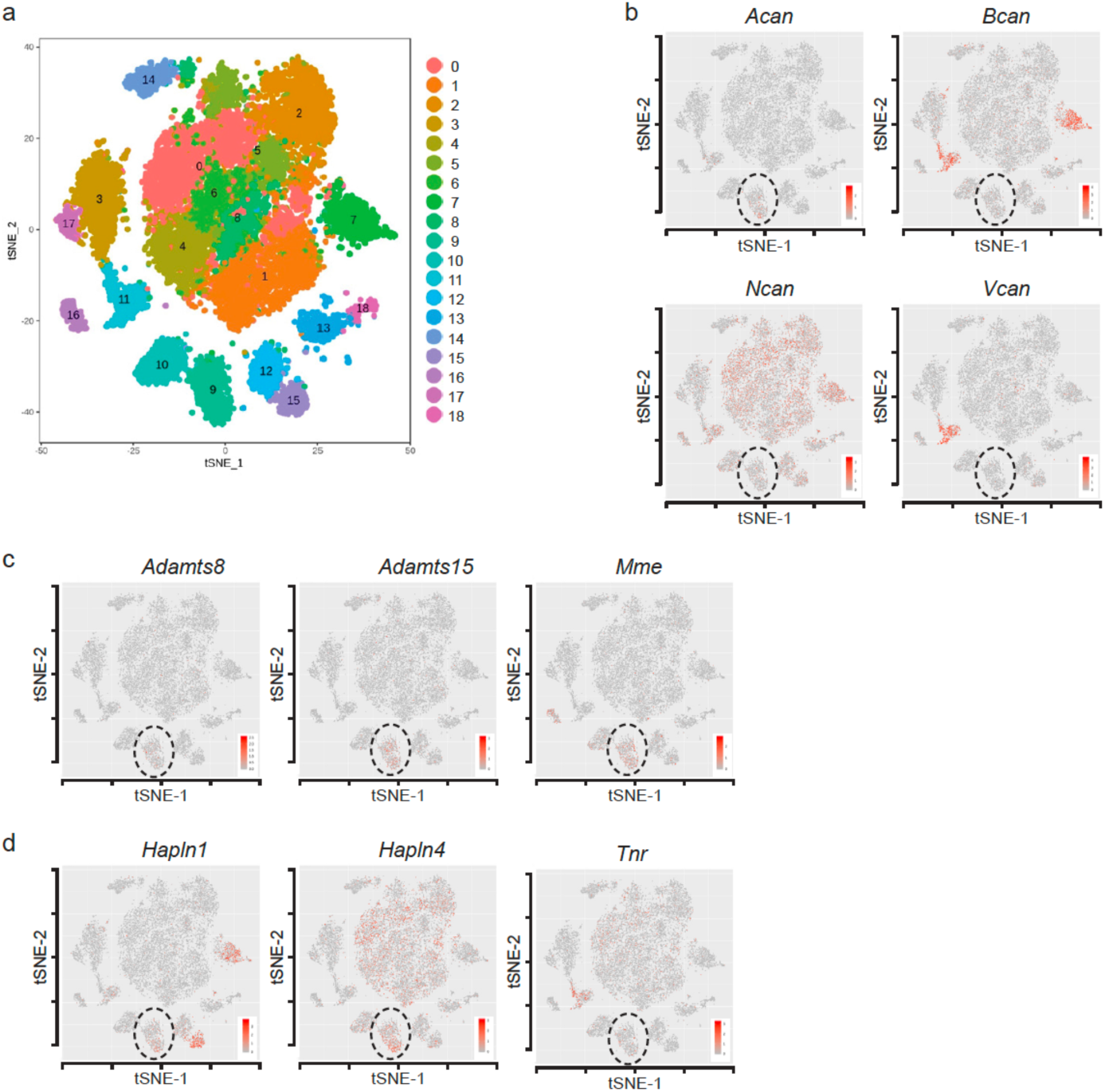
PNN components and metalloproteases are expressed by different cell types in mouse PFC. (a) Single cell RNA-sequencing (DropSeq) of PFC cells of P40-43 wild-type mice (n=8 mice). t-SNE visualization of identified cell types; cluster 9 refers to PV^+^ cells. (b-d) Expression localisation of critical components of PNN structure and remodeling, including lecticans (b, *Acan, Bcan, Ncan*, and *Vcan*), metalloproteases known to be expressed by PV^+^ cells (c, *Adamts8, Adamts15*, and *Mme*) and other critical PNN components (d, *Halpn1, Halpn4*, and *Tnr*).

Next, we analyzed which cell types express mRNAs coding for known PNN structural components and remodelling enzymes (metalloproteases). We confirmed that the metalloproteases *Adamts8, Adamts15*, and *Mme* were highly enriched in PV^+^ cells, (Fig. 3c and Supplementary Fig. 5), as previously reported^40,41^. Of the 4 analyzed lecticans, *Neurocan* was the most abundant and was expressed by different neuronal cell types and astrocytes. *Brevican* was also expressed by different cell types, including PV^+^ cells, but it was highly enriched in astrocytes and in CPSG4-expressing NG2^+^ polydendrocytes. The latter were also the major source of *Versican* (Fig. 3b and Supplementary Fig. 5). *Acan*, on the other hand, was mostly expressed by PV^+^ cells, and to a lesser extent, by polydendrocytes (Fig. 3b and Supplementary Fig. 5). None of the other PNN components analyzed (*Hapln1, Hapln4, Tnr*) showed cell type-specific expression (Fig. 3d and Supplementary Fig. 5).

Aggrecan, coded by *Acan*, plays a critical role in PNN formation, as shown in brain-wide homozygous *Acan* mutant mice and by virally-induced *Acan* deletion^27^. To investigate whether *Acan* expression is regulated by Hdac2 in PV^+^ cells, we analyzed *Acan* transcripts by RNAscope hybridization *in situ* in PFC, BLA and SSCX of adult *PV-Cre*; *Hdac2*^*lox/lox*^ mice and control littermates (Fig. 4). We confirmed that a majority of PFC PV^+^ cells expressed *Acan* in all the three regions in wildtype mice (percentage of *PV*^*+*^*Acan*^*+*^/*PV*^*+*^ cells = 71.4 ± 3.6 %, n=7 mice). Interestingly, the number of *PV*^*+*^ mRNA puncta in PV^+^ cells expressing *Acan* was significantly higher than in PV^+^ cells where *Acan* mRNA could not be detected (Supplemental Figure 6). Since low expressing PV cells have been suggested to be more plastic^17^, it is possible that *Acan* levels may be inversely correlated with PV cells’ potential for plasticity. In *PV-Cre*; *Hdac2*^*lox/lox*^ mice, *Acan* expression was significantly reduced specifically in the PFC, but not in the SSCX or the BLA, compared to control littermates (Fig. 4a, b). This decrease in *Acan* expression was accompanied by a significant decrease in the percentage of PV^+^ cells surrounded by Aggrecan immunostaining specifically in the PFC (Fig. 4c, d). Unaltered Aggrecan expression levels in SSCX of mutant mice is consistent with the observation that the percentage of PNN^+^PV^+^ cells in the SSCX was not affected (Fig. 2 a, b). Conversely, the percentage of PNN^+^PV^+^ cells was reduced in the BLA in absence of parallel changes in the percentage of Aggrecan^+^PV^+^ cells. Taken together, these data strongly point towards a strict requirement for Aggrecan in PNN formation specifically around cortical PV^+^ cells.

**Figure 4.**
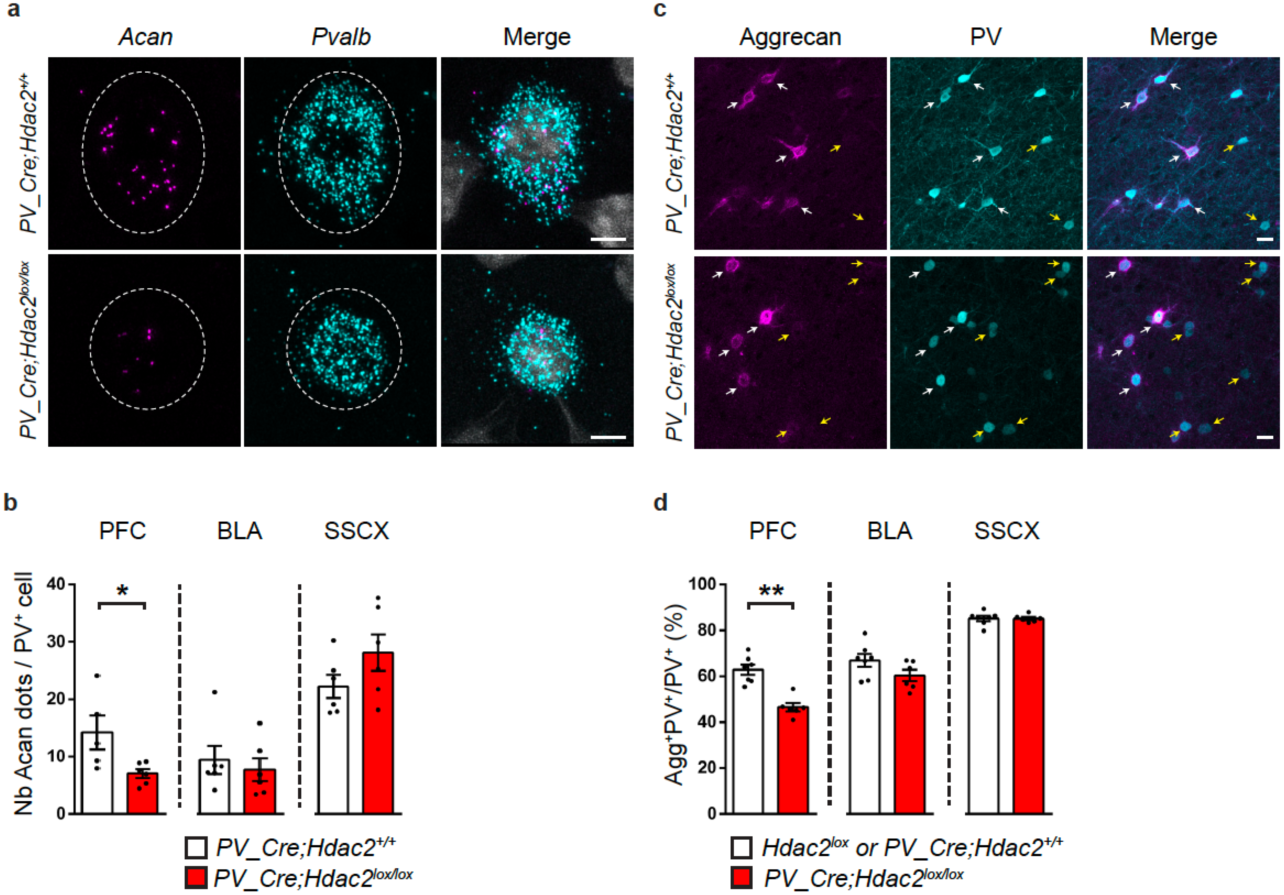
PV^+^ cell-specific *Hdac2* deletion leads to reduced *Acan* mRNA expression and Aggrecan condensation around PFC PV^+^ cell somata. (a) Representative images of fluorescent RNAscope *in situ* hybridization against *Acan* (Aggrecan, magenta) and *Pvalb* (PV, cyan) in prefrontal cortex of *PV-Cre;Hdac2*^*lox/lox*^ and control littermates (DAPI labeling is in grey). Scale bar, 5 µm. (**b**) The mean number of *Acan* dots detected in *Pvalb*^+^ cells is significantly lower in the PFC, but not in BLA or SSCX, of conditional knockout mice compared to their control littermates. PFC: Mann-Whitney test, *P*=0.0173, *PV-Cre;Hdac2*^*+/+*^n=5, *PV-Cre;Hdac2*^*lox/lox*^ n=6. BLA: Mann-Whitney test, *P*=0.3874, *PV-Cre;Hdac2*^*+/+*^ n=6, *PV-Cre;Hdac2*^*lox/lox*^ n=6. SSCX: Mann-Whitney test, *P*=0.1797, *PV-Cre;Hdac2*^*+/+*^ n=6, *PV-Cre;Hdac2*^*lox/lox*^ n=6. (**c**) Prefrontal cortex coronal sections labelled for Aggrecan (cyan) and PV (magenta) in *PV-Cre;Hdac2*^*lox/lox*^ and control littermates. White arrows indicate PV^+^Aggrecan^+^ cell bodies, while yellow arrows point to PV^+^Aggrecan^-^ cell bodies. Scale bar, 10 µm. (**d**) The percentage of PV^+^ cell bodies surrounded by aggrecan is significantly reduced in the PFC, but not BLA or SSCX, of conditional knockout mice compared to their control littermates. PFC: Mann Whitney test, *P*=0.0012, BLA: Mann Whitney test *P*=0.0513, SSCX: Mann Whitney test *P*=0.6037. For all three brain regions, *Hdac2*^*lox/lox*^ or *PV-Cre;Hdac2*^*+/+*^n=7; *PV-Cre;Hdac2*^*lox/lox*^ n= 6. Data represent mean ± s.e.m. Black circle represent individual data points. * P < 0.05, ** *P* < 0.01.

The effects of PV^+^ cell-specific *Hdac2* deletion on PNN aggregation can be due to a direct regulation of the expression of critical PNN components by Hdac2, or due to secondary homeostatic effects caused by long-term *Hdac2* deletion. To test whether Hdac2 regulates Aggrecan expression in PV^+^ cells, we virally re-expressed Hdac2 with EGFP or EGFP alone specifically in PFC PV^+^ cells of adult *PV-Cre*; *Hdac2*^*lox/lox*^ and *PV_Cre*; *Hdac2*^*+/+*^ mice for three weeks using an inverted double-floxed *Hdac2* coding region driven by a ubiquitous EF-1α promoter (Fig. 5a). Consistent with our previous data (Fig. 4d), we observed a significant reduction in the percentage of PV^+^ cells enwrapped by Aggrecan in conditional KO compared to wildtype littermates when both were injected with the control virus (AAV_DIO_EGFP; Fig. 5b, c). On the other hand, injection of AAV_DIO_Hdac2-T2A-EGFP was sufficient to rescue the percentage of Aggrecan^+^ PV^+^ cells (Fig. 5c) and to increase (albeit not significantly) the number of PV^+^ cells surrounded by WFA-stained PNN in *PV-Cre*; *Hdac2*^*lox/lox*^ mice (Fig. 5d, e).

**Figure 5.**
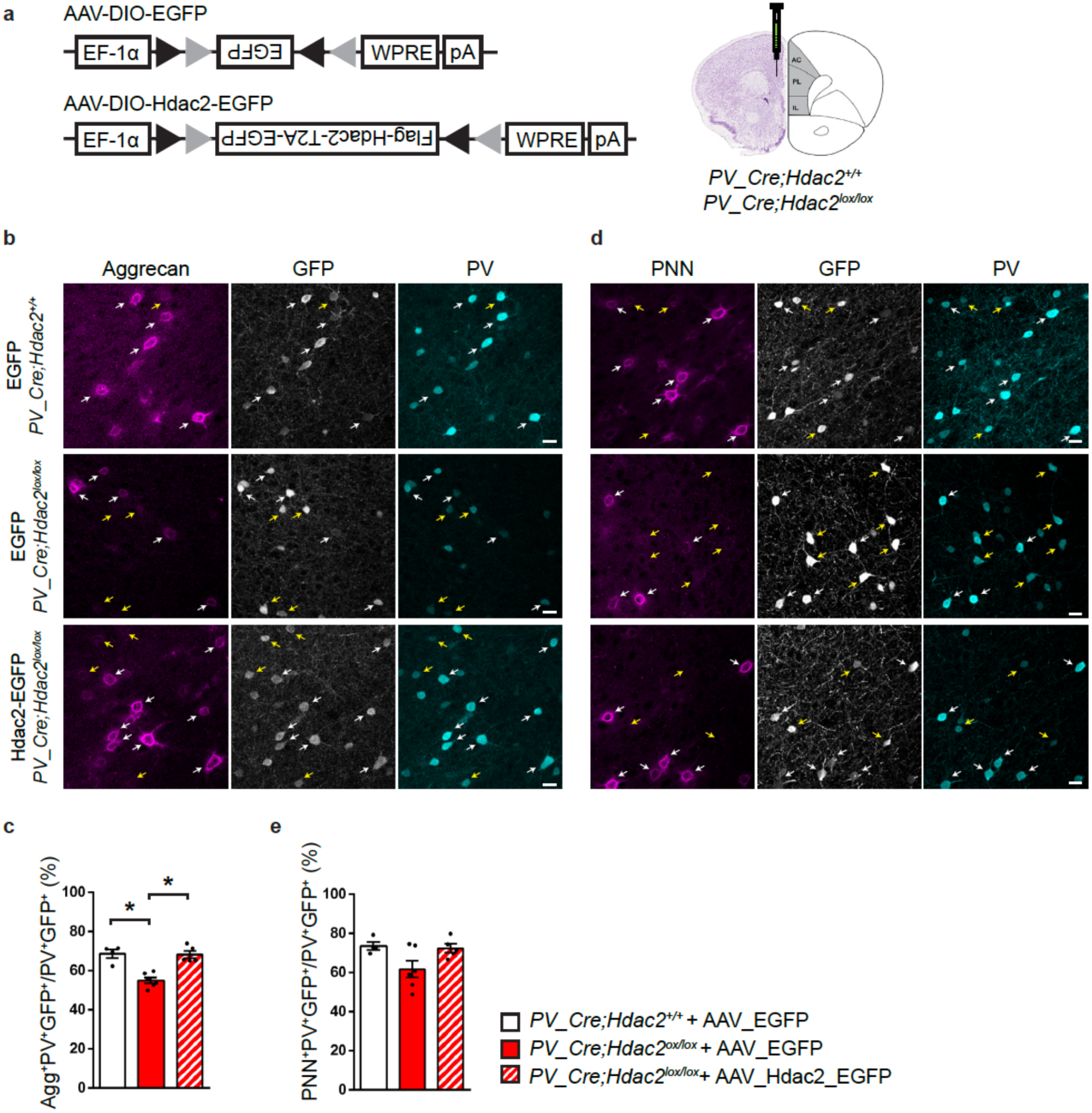
PV^+^ cell-specific Hdac2 re-expression in *PV_Cre;Hdac2*^*lox/lox*^ mice rescues Aggrecan condensation around PFC PV^+^ cell somata. (a) Schematic representation of Cre-dependent viral vectors and experimental approach. (b) Immunolabelling of Aggrecan (magenta), GFP (grey) and PV (cyan) in PFC coronal sections from *PV-Cre;Hdac2*^*+/+*^ and *PV-Cre;Hdac2*^*lox/lox*^ mice injected with the control AAV (AAV_DIO_EGFP) and from *PV-Cre;Hdac2*^*lox/lox*^ mice injected with the Cre-dependent Hdac2 expressing virus (AAV_DIO_Hdac2_T2A_EGFP). White arrows point to PV^+^GFP^+^Aggrecan^+^ cell bodies, while yellow arrows indicate PV^+^GFP^+^Aggrecan^-^ cell bodies. Scale bar, 10 µm. (**c**) Percentage of PV^+^GFP^+^ cell bodies surrounded by Aggrecan. Kruskall-Wallis with Dunn’s *posthoc* test, *P*=0.0008. (**d**) Labelling for WFA (PNN, magenta), GFP (grey) and PV (cyan) in the three experimental groups. White arrows indicate PV^+^GFP^+^PNN^+^ cell bodies, while yellow arrows indicate PV^+^GFP^+^PNN^-^ cell bodies. Scale bar, 10 µm. (**e**) Percentage of PV^+^GFP^+^ cell bodies surrounded by WFA-stained PNN in the prefrontal cortex. Kruskall-Wallis with Dunn’s *posthoc* test, *P*=0.2603. Number of mice: *PV-Cre;Hdac2*^*+/+*^ + AAV_DIO_EGFP, n=4, *PV-Cre;Hdac2*^*lox/lox*^ + AAV_DIO_EGFP, n=6, *PV-Cre;Hdac2*^*lox/lox*^ + AAV_DIO_Hdac2_T2A_EGFP, n= 5. Data represent mean ± s.e.m. Black circle represent individual data points. * *P* < 0.05.

To further support the hypothesis that Hdac2 activity in PV^+^ cells regulates Aggrecan levels, we explored the effects of acute Hdac2 pharmacological inhibition. We found a significant decrease in the number of *Acan* mRNA molecules in PFC PV^+^ cell somata (Fig. 6a, b) and in the proportion of PV^+^ cells enwrapped by Aggrecan^+^ nets (Fig. 6d, e) 17h after BRD6688 i.p. injection in wild-type mice. Conversely, the same treatment was unable to further reduce *Acan*/Aggrecan expression in *PV-Cre; Hdac2*^*lox/lox*^ mice (Fig. 6c, f).

**Figure 6.**
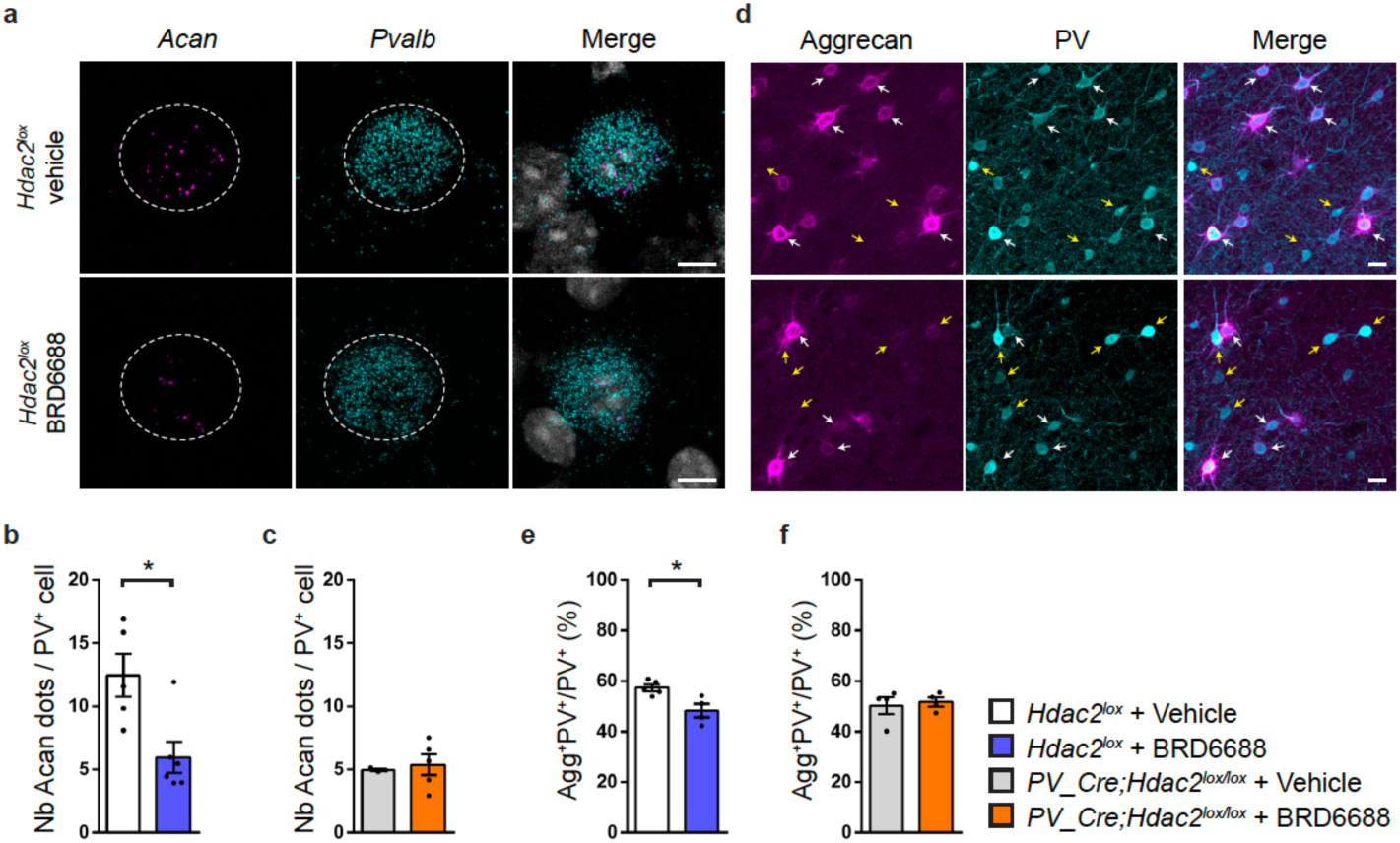
Hdac2 inhibition decreases *Acan* mRNA expression and Aggrecan agglomeration around PFC PV^+^ cell somata in wild-type but not in *PV-Cre;Hdac2*^*lox/lox*^ mice. (a) Representative images of fluorescent RNAscope *in situ* against *Acan* (Aggrecan, magenta) and *Pvalb* (PV, cyan) around a DAPI (grey) nucleus in PFC of *Hdac2*^*lox/lox*^ mice 17 hours after i.p. injection of either the Hdac2 inhibitor BRD6688 or the vehicle. Scale bar, 5 µm. (b) The mean number of *Acan* dots present around a *Pvalb*^+^ cell is reduced in BRD6688-treated wild-type (Mann-Whitney test, *P*=0.0303) (c) but not conditional knockout mice (Mann-Whitney test, *P*=0.6786). Number of *Hdac2*^*lox/lox*^ injected with vehicle, n=5 or BRD6688, n=6. Number of *PV-Cre;Hdac2*^*lox/lox*^ mice injected with vehicle n=3 or BRD6688, n=5. (d) PFC coronal sections immunolabeled for Aggrecan (cyan) and PV (magenta) in *Hdac2*^*lox/lox*^ mice 17 hours after i.p. injection of either the Hdac2 inhibitor BRD6688 or the vehicle. Scale bar, 10 µm. (e) The percentage of PV^+^ cell bodies surrounded by Aggrecan is reduced by BDR6688 injection in control (Mann-Whitney test, *P*=0.0317) (f) but not conditional knockout mice (Mann-Whitney test, *P*=0.8286). Number of *Hdac2*^*lox/lox*^ injected with vehicle, n=5 or BRD6688, n=4. Number of *PV-Cre;Hdac2*^*lox/lox*^ mice injected with vehicle, n=4 or BRD6688, n=4. Graph bars represent mean ± s.e.m. Circles represent individual mouse values. * *P* < 0.05.

Overall, these results show that PV^+^ cells are the major source of Aggrecan in the PFC and that Hdac2 modulates PNN aggregation by regulating *Acan* transcription in PV^+^ cells.

### Reduced *Hdac2* activity in PFC PV^+^ cells promotes dynamic changes in *Acan* expression following extinction training

Class 1 Hdac (including Hdac2) inhibition has been shown to prime the expression of neuroplasticity-related genes^22^. We thus asked whether *Acan* expression might be more dynamic in *Hdac2*^*−/−*^ PV^+^ cells following extinction training. To answer this question, we collected brains from *PV-Cre; Hdac2*^*lox/lox*^ mice and control littermates 3 hours after the end of extinction training (Fig. 7a, b) and quantified *Acan* mRNA expression in PV^+^ cells from the PFC (Fig. 7c, d). We found that, although *Acan* expression was reduced in PFC PV^+^ cells from naive mutant mice compared to wild-type littermates (Fig. 4a, b), its expression reached wild-type levels after extinction training (Fig. 7c, d). Next, we asked whether acute pharmacological Hdac2 inhibition could promote *Acan* dynamic changes triggered by extinction learning. While naive mice injected with BRD6688 showed reduced *Acan* levels (Fig. 6a, b), mice injected with BRD6688 following fear learning but before the onset of extinction training showed no significant difference between *Acan* levels compared to vehicle-treated controls 3 hours after the end of extinction training (Fig. 7g, h).

**Figure 7.**
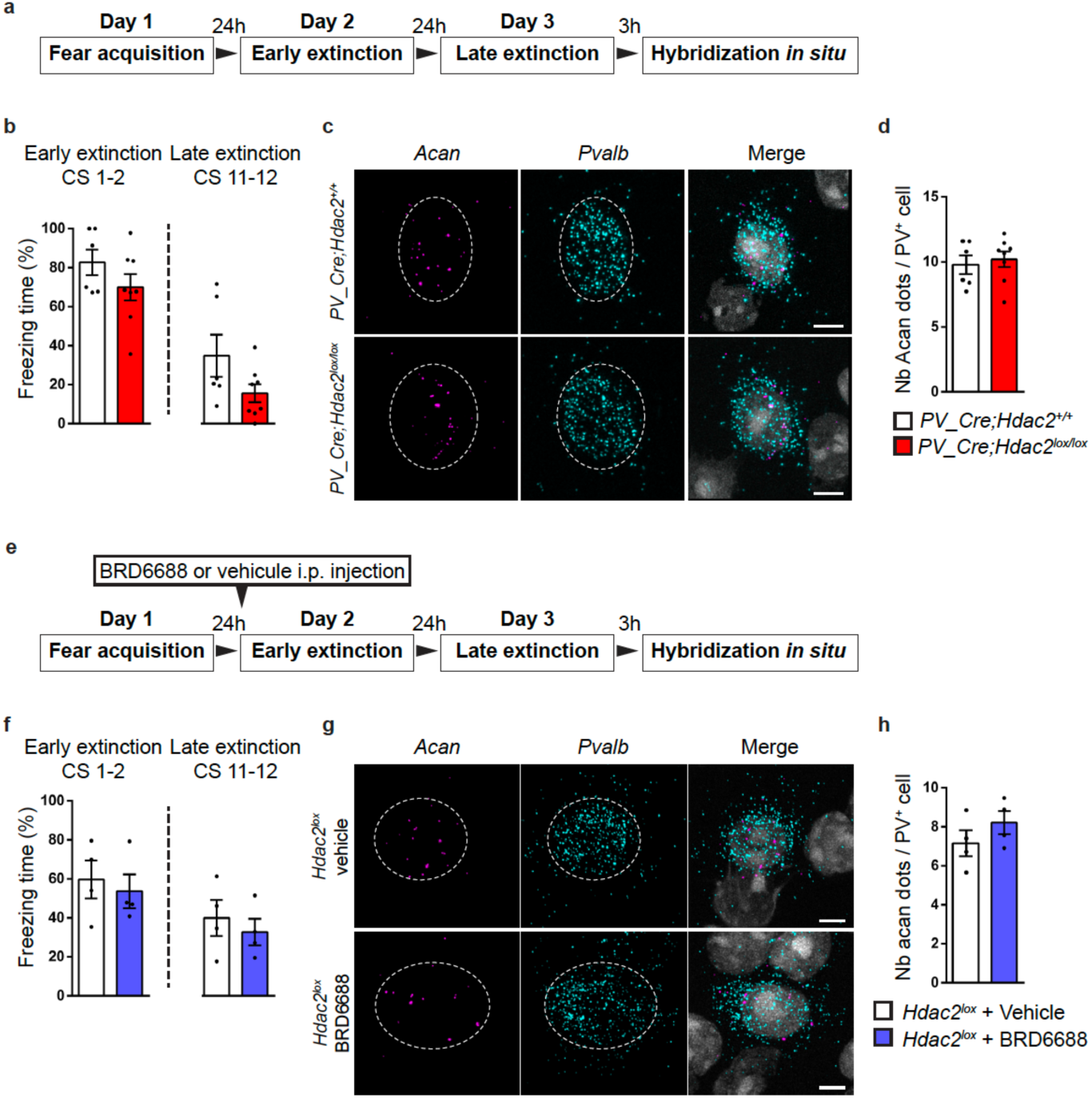
Hdac2 deletion or pharmacological inhibition promotes dynamic changes in *Acan* expression in PFC PV^+^ cells following fear extinction training. (a) Schematic representation of the experimental protocol. Brains of *PV-Cre;Hdac2*^*lox/lox*^ mice and their control littermates have been collected 3 hours after late extinction training for *in situ* hybridization. (b) Both *PV-Cre;Hdac2*^*lox/lox*^ mice and controls show efficient fear extinction. Repeated two-way ANOVA; F_genotype_ (1,12)=7.554, *P*=0.0177, F_extinction_(11,132)=9.148, *P*<0.0001, F_genotype_*_extinction_(11,132)=1.107, *P*=0.3605. (c) Representative images of fluorescent RNAscope *in situ* against *Acan* (Aggrecan, magenta) and *Pvalb* (PV, cyan) around a DAPI (grey) nucleus in PFC of *PV-Cre;Hdac2*^*+/+*^ and *PV-Cre;Hdac2*^*lox/lox*^ mice 3 hours after late extinction training. Scale bar, 5 µm. (d) The mean number of *Acan* dots present around a prefrontal *Pvalb*^+^ cell 3 hours after late extinction training is comparable between *PV-Cre;Hdac2*^*+/+*^ and *PV-Cre;Hdac2*^*lox/lox*^ mice (Mann-Whitney test, *P*=0.8112). Number of mice: *PV-Cre;Hdac2*^*+/+*^ n=6; *PV-Cre;Hdac2*^*lox/lox*^ n=8. (e) Schematic representation of the experimental protocol. Brains of BRD6688-injected or vehicle-injected mice 6 hours before extinction training have been collected 3 hours after late extinction training for *in situ* hybridization. (f) Both BRD6688-injected and vehicle-injected controls show efficient fear extinction. Repeated two-way ANOVA; F_treatment_ (1,6)=1.093, *P*=0.3362, F_extinction_(11,66)=3.902, *P*=0.0002, F_treatment_*_extinction_ (11,66)=0.7446, *P*=0.6922. (g) Representative images of fluorescent RNAscope *in situ* against *Acan* (Aggrecan, magenta) and *Pvalb* (PV, cyan) around a DAPI (grey) nucleus in PFC of BRD6688-injected and vehicle-injected mice 3 hours after late extinction training. Scale bar, 5 µm. (h) The mean number of *Acan* dots present around a prefrontal *Pvalb*^+^ cell 3 hours after late extinction training is comparable between BRD6688-injected and vehicle-injected mice (Mann-Whitney test, *P*=0.2). Number of mice: vehicle-injected n=4; BRD6688-injected n=4.

Altogether, these results indicate that either PV^+^ cell-specific Hdac2 deletion or pharmacological inhibition allows for dynamic changes in *Acan* expression following extinction training.

### *Acan* knock-down during extinction training is sufficient to limit the spontaneous recovery of fear memories with time

Finally, we asked whether the observed dynamic changes in *Acan* levels caused by Hdac2 inhibition were sufficient to promote enhanced retention of extinction memory over time. To answer this question, we sought to transiently decrease *Acan* expression after fear learning but before extinction training. To this purpose, we used a chimeric fusion protein comprising a double-stranded RNA binding domain fused to a brain targeting peptide, namely TARBP-BTP. to target siRNA to the brain ^42,43^. Previous data showed that using TARBP-BTP fusion protein as a carrier facilitated siRNA delivery across the brain-blood barrier to the brain^42^. We observed a significantly reduced number of *Acan* mRNA molecules in PFC PV^+^ cells compared to the control group 48hrs after TARBP-BTP/*Acan*-siRNA intravenous injection in wild-type adult mice (Fig. 8 a-c), demonstrating that siRNAs were efficiently delivered to the brain parenchyma and could enter PV^+^ cells. We then injected either TARBP-BTP/*Acan-*siRNA or TARBP-BTP/*CtrlNeg*-siRNA in wild-type mice, 24hrs after fear learning, and then subjected them to fear extinction training 24hrs after injection (Fig. 8d). Compared to control TARBP-BTP/*CtrlNeg*-siRNA mice, TARBP-BTP/*Acan-* siRNA-injected mice presented significantly less freezing behavior during the fear retrieval test (Fig. 8e).

**Figure 8.**
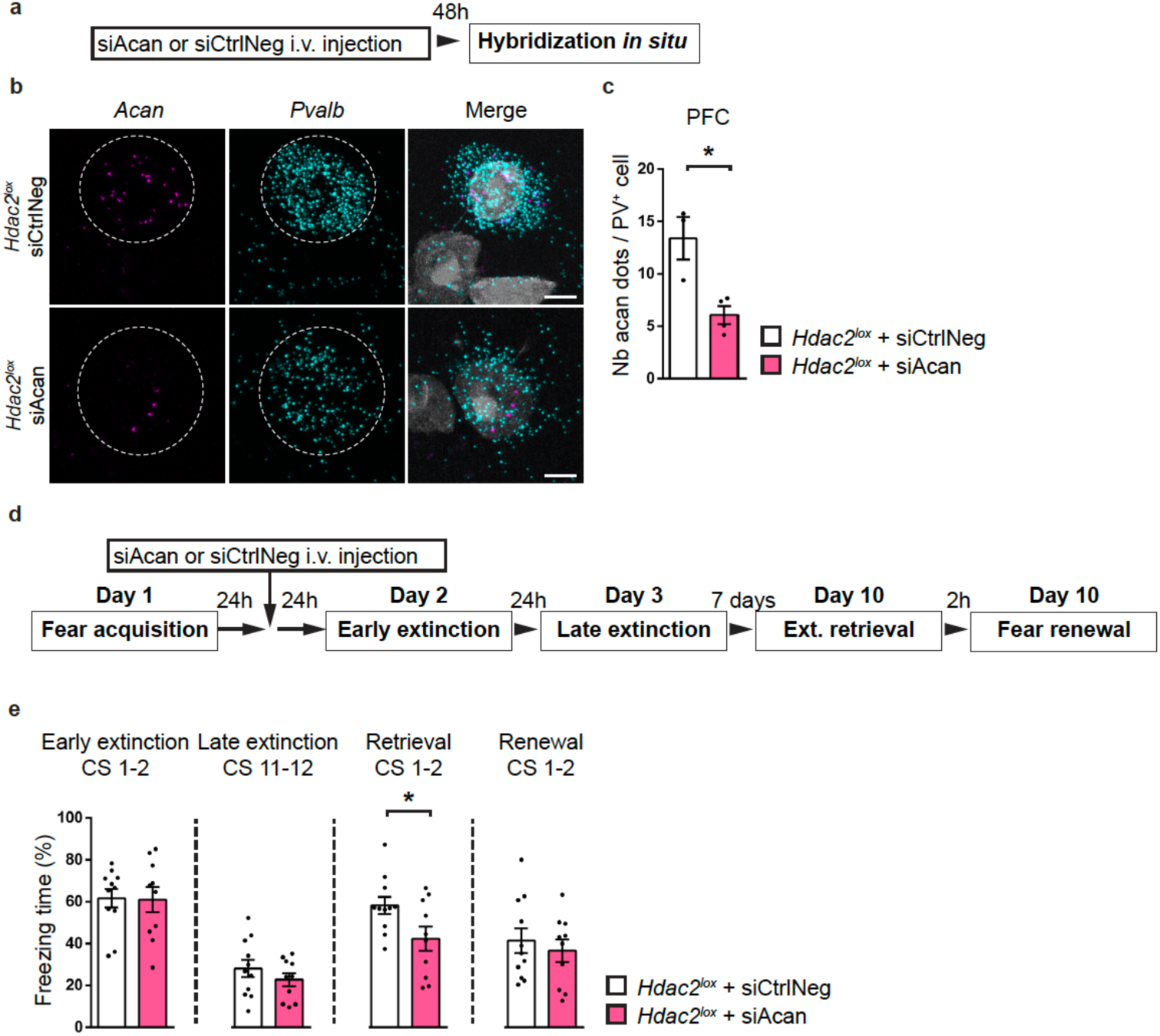
*Acan* knock-down during extinction training is sufficient to increase the persistence of extinction memories. (a) Schematic representation of the experimental protocol. Brains of adult *Hdac2*^*lox*^ mice have been collected 48h after the intravenous injection (i.v.) of either TARBP-BTP:siCtrlNeg or TARBP-BTP:siAcan. (b) Representative images of fluorescent RNAscope *in situ* against *Acan* (Aggrecan, magenta) and *Pvalb* (PV, cyan) around a DAPI (grey) nucleus in PFC of *Hdac2*^*lox*^ mice 48 hours after the i.v. injection described in (a) and quantified in (c). Scale bar, 5 µm. (c) The mean number of *Acan* dots present around a prefrontal *Pvalb*^+^ cell 48 hours after the i.v. injection of TARBP-BTP:siAcan is significantly decreased compare to TARBP-BTP:siCtrlNeg (Unpaired two tailed t-test, P=0.0139). Number of mice: *Hdac2*^*lox*^ + TARBP-BTP:siCtrlNeg n=3; *Hdac2*^*lox*^ + TARBP-BTP:siAcan n=4. (d) Schematic representation of the experimental protocol. Ext.: extinction. (e) TARBP-BTP:siCtrlNeg-injected and TARBP-BTP:siAcan-injected mice show efficient fear extinction. Repeated two-way ANOVA; F_treatment_ (1,19)=1.109, *P*=0.3054, F_extinction_(11,209)=18.85, *P*<0.0001, F_treatment_*_extinction_ (11,209)=1.052, *P*=0.4021. One week after extinction training, freezing levels are significantly different in the spontaneous recovery test between TARBP-BTP:siCtrlNeg- and TARBP-BTP:siAcan-injected *Hdac2*^*lox*^. Extinction retrieval unpaired two-tailed *t*-test, *P*=0.0349; fear renewal, unpaired two-tailed *t*-test, *P*=0.5608. Number of *Hdac2*^*lox*^ mice injected with TARBP-BTP:siCtrlNeg, n=11 or TARBP-BTP-siAcan, n=10. Graph bars represent mean ± s.e.m. Circles represent individual mouse values. * *P* < 0.05.

All together these data showed that *Acan* knock-down in PV^+^ cells prior to extinction training is sufficient to render extinction memories stronger and limits the spontaneous recovery of fear memories with time.

## Discussion

The maturation of GABAergic circuits, in particular of PV^+^ cells, is one of the factors that restricts critical period plasticity in the adult brain^11,13,36^. Hence, controlled manipulation of cortical PV^+^ cell function can reinstate heightened plasticity, creating an opportunity for therapeutic intervention^44,45^. To this purpose, it is essential to gain a better understanding of the molecular mechanisms determining PV^+^ cell maturation state in the adult brain. Here, we showed that the epigenetic landscape of PV^+^ cells regulates their plastic potential, affecting adult neuroplasticity. In particular, our results demonstrate that PV^+^ cell-specific *Hdac2* deletion reduces PNN aggregation around PV^+^ cell somata and PV^+^ cell synapse density in naive mice, while promoting PV^+^ cell synaptic remodeling following extinction training. These anatomical changes are accompanied by decreased spontaneous recovery of fear memories with time in mutant mice. We further found that Hdac2 levels controlled the expression of the lectican Aggrecan, a critical regulator of PNN aggregation which in turn limits adult neuroplasticity, by regulating its transcription specifically in PFC PV^+^ cells. Finally, siRNA-mediated brain-targeted *Acan* reduction after fear acquisition coupled with extinction training was sufficient to reduce the spontaneous recovery of fear memory with time in adult wild-type mice, indicating a potential therapeutic target for enhancing extinction-dependent plasticity.

A previous study by Gogolla and collaborators^28^ showed that degradation of PNNs by chondroitinase ABC injection in adult BLA rendered subsequently acquired fear memories susceptible to erasure following extinction learning. However, the same treatment had no effect when applied after fear learning, but before extinction training. In contrast, our data showed that global *Acan* reduction after fear learning was sufficient to reduce the spontaneous recovery of fear memories one week after extinction training. It is possible that PNN remodeling in PFC and not, or not exclusively, in the BLA, is essential for long term retention of extinction learning. Alternatively, the dynamic regulation of PNN expression, more than its absolute levels, may play a key role in this process. Indeed, a transiently decrease in *Acan* expression after fear learning but before extinction training promoted increased retention of extinction memories, suggesting that dynamic changes of PNNs could be a critical step in this process. To test this hypothesis, we choose to deliver siRNA against *Acan* using a peptidic carrier that crosses the BBB. Weather transient *Acan* changes and PNN remodeling selectively in PFC are sufficient to affect the spontaneous recovery of fear memories, or whether PNN remodeling needs to occur in the BLA too, remain to be established.

Numerous studies have addressed the role of epigenetic modifiers, through pharmacological inhibition or genetic deletion of different HDACs, on fear memory formation and extinction^46^. In particular, an elegant study demonstrated that pharmacological inhibition of Hdac2 during contextual fear extinction prevents the spontaneous recovery of freezing behaviour during the recall of remote fear memories by upregulating neuroplasticity-related gene expression^22^. However, whether Hdac2 plays a different role in different neuron types thus leading to distinct behavioural outputs, is not clear. A previous study showed that *Hdac2* deletion in post-mitotic forebrain glutamatergic neurons (using CaMKII-Cre mice) led to a robust acceleration of extinction learning, which was however followed by a normal spontaneous recovery of fear memories 5 days after the end of the extinction^25^. Conversely, our data showed that *Hdac2* deletion specifically in postnatal PV^+^ cells did not affect the rate of extinction learning but led to decreased spontaneous recovery of fear memories with time. The specific role of Hdac2 expressed by PV^+^ cells in limiting the long-term retention of extinction memories was further supported by the fact that the beneficial effect of Hdac2 inhibitor and PV^+^ cell-specific deletion of *Hdac2* on preventing the recovery of fear memory were not additive. These different results could be explained by partly different roles played by glutamatergic and PV^+^ neurons in fear extinction learning and retention. It is also conceivable that HDAC2 regulates a different set of genes in different cell types, thus leading to different cellular responses.

The main problem with fear extinction is the reappearance, or spontaneous recovery, of the original fear after extinction training. To explain this phenomena, two opposite models of extinction have been proposed and strongly debated: one posits that the extinction training leads to the erasure of the fear memory itself^16,28^, while the second proposes that extinction creates new inhibitory learning, which controls the fear memory^47,48^. It has been suggested that extinction learning could be seen as an equilibrium between erasure and inhibition, where any behavioural manipulation which influences fear expression would either promote a functional erasure of the original fear trace or have permissive effects on the new inhibitory memory^49^. In the field of cognitive neuroepigenetics, it has been proposed that “there exists an equilibrium in epigenetic states that can be pushed in one direction or another” ^49^. The data presented in this work support the hypothesis that increasing chromatin acetylation, likely at specific genes, enhances fear extinction memory and its retention over time. In particular, we identified PV^+^ cell-epigenetic state as a major determinant in the long-term retention of extinction memories. Whether Hdac2 inhibition in PV^+^ cells leads to stronger inhibitory memory opposing the fear memory, or towards erasure of the original fear memory is not clear. However, the lack of generalisation of fear extinction memory, as observed by similar freezing times for the renewal test in *PV-Cre;Hdac2*^*lox/lox*^ versus *PV-Cre;Hdac2*^*+/+*^ mice (Suppl. Fig. 2) or following Hdac2 inhibition after fear memory acquisition (Fig. 1g), argues against the erasure of the original fear memory, instead supporting the hypothesis that PV^+^ cell heightened plastic remodelling could contribute to forming stronger (or more stable) inhibitory memory^15^.

The development of cortical PV^+^ cells is a prolonged process which plateaus only by the end of adolescence^35^. During the postnatal maturation phase, cortical PV^+^ cells form exuberant innervation fields characterized by baskets of perisomatic synapses. In addition, during their maturation phase, PV^+^ cell somata and dendrites are progressively enwrapped in PNNs, thought to stabilize their synaptic inputs^18,50^ and protect them from oxidative stress^51^. Our data suggest that PV^+^ cell-epigenetic modifications likely regulate their maturation state, since *PV-Cre;Hdac2*^*lox/lox*^ mice showed significantly reduced PNN condensation around PV^+^ cells and decreased density of PV^+^ cell perisomatic synapses in the PFC and BLA, two brain regions implicated in fear behaviour regulation. Our data is consistent with the observation that PV^+^ cell-restricted *Hdac2* deletion reduces inhibitory input onto pyramidal cells in the visual cortex of adult mice, and coincides with enhanced long-term depression that is more typical of younger mice^52^. Increased acetylation levels in PV^+^ cells likely modulate their plasticity potential as well, since we observed increased *Acan* expression and synaptic remodelling following extinction training. Such heightened synaptic remodelling and decreased expression of critical PNN components might contributes to long-term retention of extinction memory. In accordance with this hypothesis, Chen et al.^14^ showed that extinction learning increased the fraction of PV^+^ cells expressing low PV levels, which are thought to be more plastic^17^. They further showed that chemogenetic-mediated manipulations of PV^+^ cell activity affected mouse freezing during extinction learning, with more active PV^+^ cells leading to higher freezing. However, this study did not explore whether and how these manipulations affected long-term spontaneous recovery of fear memories. Further, Trouche et al.^15^ showed that extinction training induced structural remodelling of perisomatic PV^+^ cell synapses specifically around excitatory neurons that had been previously activated during the encoding of the original fear memory (fear neurons) in the BLA. In addition, Bhagat et al.^16^ reported that PV^+^ cell-specific deletion of Nogo Receptor 1, a neuronal receptor for myelin-associated growth inhibitors, enhances both BLA PV^+^ cell synaptic structural remodelling and the retention of fear memory extinction. In *PV-Cre;Hdac2*^*lox/lox*^ mice, we observed increased remodelling of PV^+^ cell synapses in both BLA and PFC following extinction training. Elegant work demonstrated that PFC PV^+^ cell firing inversely correlates with freezing behaviour and that their optogenetic activation reduces freezing behaviour^53^, thus, it is possible that reinforcing PFC PV^+^ cell-mediated inhibitory drive by the end of extinction training could contribute to the maintenance of the extinction memory, thereby leading to reduced fear response freezing at the retrieval test.

How and to what extent PV^+^ cells control the development, integrity and dynamic of the PNNs surrounding their somata and dendrites, in a cell-autonomous fashion, is still an open question. By using single-cell transcriptomic analysis, we found that PFC PV^+^ cells express a variety of PNN molecular components and metalloproteases. While most of these factors were expressed by other cell types as well, *Acan* transcription appeared to be mostly restricted to PV^+^ cells, consistent with previous studies^54,55^. *In vitro*^56,57^ and *in vivo*^27^ experiments revealed that Aggrecan, coded by *Acan*, plays a critical role in PNN formation. For instance, brain-wide targeting of *Acan* (*Nes_Cre; Acan*^*lox/lox*^ mice) or virally-mediated local *Acan* deletion in adult visual cortex led to complete loss of WFA-stained PNN, while conditional deletion of one *Acan* allele was sufficient to reduce WFA staining intensity^27^. Therefore, the reduced Aggrecan levels we observed in *PV_Cre;Hdac2*^*lox/lox*^ or BRD6688-injected mice likely contributes to impaired aggregation of other extracellular matrix molecules into PNNs. Our data showed that PV^+^ cell-specific *Hdac2* deletion reduced *Acan/*Aggrecan expression levels in PV^+^ cells, while re-introduction of Hdac2 in *Hdac2*^*−/−*^ mutant cells rescued *Acan/*Aggrecan expression levels, thus suggesting that Hdac2 controls *Acan* levels in PV^+^ cells. By determining Aggrecan expression levels, PV^+^ cells may cell-autonomously modulate PNN dynamics and integrity, thereby regulating the stability of their synaptic inputs^18,50,58^. Further, a recent study showed that 4 days of monocular deprivation was sufficient to produce a strong shift in ocular dominance in adult *Acan*^*lox/lox*^ mice injected with AAV-Cre, suggesting that *Acan* expression in the adult brain acts as a brake on neuroplasticity^27^. Decreased *Acan* levels in *PV-Cre;Hdac2*^*lox/lox*^ or BRD6688-injected mice can therefore increase neuroplasticity leading to higher retention of extinction memory. Consistent with this hypothesis, our data show that a brief *Acan* reduction coupled with extinction training was sufficient to decrease the spontaneous recovery of fear memory over time. Furthermore, we showed that PV^+^ cell-specific *Hdac2* deletion reduced *Acan* transcription selectively in PFC, but not in somatosensory cortex, PV^+^ cells. This surprising regional specificity suggests that PV^+^ cell circuits in different adult cortical areas might differ more, at least in term of epigenetic regulation, than what was previously assumed. In particular, region-specific dynamic epigenetic regulation of PV^+^ cells could increase plasticity of PFC networks compared to primary sensory areas in the adult brain.

In contrast to what we observed in the PFC, the reduced numbers of PV^+^ cells enwrapped by PNN in the BLA of *PV-Cre;Hdac2*^*lox/lox*^ mice was not accompanied by reduced *Acan*/Aggrecan expression, indicating that different molecular organizers might be at play in the two regions. In BLA, PNN enwrap not only PV^+^ cells, but also a population of excitatory neurons^59^, therefore PNNs components produced by excitatory neurons may contribute to PNN aggregation around PV^+^ cells. Finally, consistent with previous findings^40^, we observed that the expression of the metalloproteases *Adamts8, Adamts15* and *Mme* was highly enriched in, but not restricted to, PV^+^ cells. Increased metalloproteases expression might lead to Aggrecan cleavage and PNN disassembly. Whether their expression levels are regulated, directly or indirectly, by *Hdac2* remains to be explored.

Finally, we observed that while *Acan* gene expression in PFC PV^+^ cells was lower in naïve *PV_Cre;Hdac2*^*lox/lox*^ and in BRD6688-injected wild-type mice compared to their respective control groups, the difference was lost after extinction training, suggesting that Hdac2 might limit dynamic changes in *Acan* expression. Increased *Acan* expression likely leads to increased PNN aggregation and stabilisation surrounding PV^+^ cells^23,45,46^. It has been suggested that new PNN formation plays a critical role in memory consolidation, by modulating PV^+^ cell activity^60^; therefore, it is possible that the ability to dynamically regulate *Acan*, and thus PNN formation/stability, and not absolute levels of *Acan*, determines the long-term retention of fear extinction memory. A recent study showed that Hdac2 S-nitrosylation was upregulated in the hippocampus during the extinction of contextual fear memories. Hdac2 S-nitrosylation caused Hdac2 dislocation from the chromatin and increased chromatin acetylation, thereby priming the expression of neuroplasticity-regulated genes (“transcriptionally permissive state”)^22^. Whether the same epigenetic mechanism is at play in PFC PV^+^ cells during fear extinction learning and whether Aggrecan expression is downstream of such regulation, remains to be investigated.

Taken together, our data supports a model in which the acetylation of PV^+^ cell chromatin primes the *Acan* gene in PFC to be dynamically expressed during extinction learning. Increased Aggrecan expression in turn promotes PNN condensation^23,45,46^, and modulates PV^+^ cell-mediated inhibition in the PFC contributing to reduced fear expression. In addition, acetylation of PV^+^ cell chromatin regulates the ability of PV^+^ cell synapses to remodel their output induced by extinction training, which could promote the selective inhibition of fear memory traces. Since spontaneous recovery of fear response following extinction training remains an important challenge in exposure therapy, any manipulation that can potentiate the plasticity of PV^+^ cell connectivity, whether it is through modulating chromatin accessibility or the expression of PNN components, could in turn foster adult brain plasticity and be of future therapeutic interest.

## Material and Methods

### Animals

*Hdac2*^*lox/lox*^ (The Jackson laboratory (JAX), *B6*.*Cg-Hdac2*^*tm1*.*1Rdp*^*/J*, 022625) mice, in which exons 5 and 6 of *Hdac2* gene, encoding the HDAC-catalytic core of the protein, are flanked by loxP sites, were crossed to the *PV-Cre* (JAX, *B6*.*129P2-Pvalb*^*tm1(cre)Arbr*^*/J*, 017320) line to generate *PV-Cre*^*+/−*^ *;Hdac2*^*lox/lox*^ (*Hdac2* cKO) mice. These lines were maintained through crosses of *PV-Cre*^*+/−*^ *;Hdac2*^*lox/+*^ females and *Hdac2*^*lox/lox*^ males to generate *PV-Cre*^*+/−*^*;Hdac2*^*lox/lox*^ and *PV-Cre*^*−/−*^ *;Hdac2*^*lox/lox*^ or *PV-Cre*^*−/−*^*;Hdac2*^*lox/+*^ control littermates, or *PV-Cre*^*+/+*^*;Hdac2*^*lox/+*^ females and *Hdac2*^*lox/+*^ males to generate *PV-Cre*^*+/−*^*;Hdac2*^*lox/lox*^ and *PV-Cre*^*+/−*^*;Hdac2*^*+/+*^ control littermates, on a mixed 129sv/C57BL/6J background. Cell specificity of Cre-mediated recombination was analyzed by breeding *PV-Cre* with *RCE*^*EGFP*^ mice (JAX, *Gt(ROSA)26Sor*^*tm1*.*1(CAG-EGFP)Fsh*^*/Mjax*, 32037). All animals were maintained under a light-dark cycle (12 h light–12 h dark) in a temperature and humidity-controlled room. Food and water were available *ad libitum*. All procedures described here had been approved by the Comité Institutionnel de Bonnes Pratiques Animales en Recherche (CIBPAR) of the Research Center of Sainte-Justine Hospital in accordance with the principles published in the Canadian Council on Animal’s Care’s (Guide to the Care and Use of Experimental Animals). Room lights were kept low during all procedures.

### Mice genotyping

DNA was extracted from mouse tails and genotyped to detect the presence of Cre alleles and Hdac2 conditional and wild-type alleles. Polymerase chain reaction (PCR) was performed using either a set of 2 separated primers (F1 5’-TGGTATGTGCATTTGGGAGA-3’ and R1 5’-ATTTCACAGCCCCAGCTAAGA-3’) to identify the *Hdac2* floxed versus the wildtype allele or a set of 3 separate primers (5’-ATTTGGGAGAAGGCCAGTTT-3’, 5’-AATTTCACAGCCCCAGCTAAG-3’ and 5′-CGAAATACCTGGGTAGATAAAGC-3′) to assure the absence of the *Hdac2* null allele: with band sizes of 720 bp for the wild-type, 560 bp for the floxed and 380 bp for the null allele. The 3 separate primers used to detect *Cre* in *PV_Cre* were: F1 (5’-CAGCCTCTGTTCCACATACACTCC-3’), F2 (5’-GCTCAGAGCCTCCATTCCCT-3’) and R1 (5’-TCACTCGAGAGTACCAAGCAGGCAGGA GATATC-3’) which generate 400 bp and 526 bp (mutant and wild-type) bands. To detect the presence of the RCE allele, 3 separate primers namely, RCE-Rosa1 (5’-CCCAAAGTCGCTCTGAGTTGTTATC-3’), RCE-Rosa2 (5’GAAGGAGCGGGAGAAATGGATATG-3,) and RCE-Cag3 (5’-CCAGGCGGGC CATTTACCGTAAG-3’) were used, which generated 350bp and 550bp bands.

### Immunohistochemistry

Mice of both sexes were anesthetized, then perfused intracardially with PBS followed by 4% (w/v) paraformaldehyde (PFA) in PBS. Intact brains were extracted and post-fixed in 4% PFA/PBS overnight at 4 °C. The tissue was then cryoprotected in 30% (w/v) sucrose (Sigma) in PBS, sectioned coronally at 40μm on a cryostat (Leica VT100) and stored as floating sections in PBS. For immunohistological analysis, brain sections were blocked in 10% normal goat serum (NGS, Invitrogen, 10000C) in PBS containing 1% (v/v) Triton X-100 for 2 h at room temperature. Primary antibodies were diluted in 5% goat serum in PBS containing 0.1% (v/v) Triton X-100 and incubated 24h-48h at 4 °C. Slices were then washed in PBS (3 × 10’), incubated in the appropriate Alexa-conjugated antibodies in 5% NGS, 0,1% Triton in PBS for 2h at room temperature, washed again in PBS (3 × 10’), and mounted in Vectashield (Vector Lab, H-1000) before imaging.

The primary antibodies used in this study and their working concentrations are as follows: rabbit monoclonal anti-Hdac2 (1:500; Abcam, 32117, RRID:AB_732777); mouse monoclonal anti-PV (1:2000; Swant, 235, RRID:AB_10000343); rabbit polyclonal anti-PV (1:4000; Swant, PV27, RRID:AB_2631173); guinea pig polyclonal anti-PV (1:1000; Synaptic Systems, 195004, RRID:AB_2156476); mouse monoclonal anti-gephyrin (1:500; Synaptic Systems, 147021, RRID:AB_22325461); rabbit polyclonal anti-aggrecan (1:500; Millipore, AB1031, RRID:AB_90460); chicken polyclonal anti-GFP (1:1000; Abcam, 13970, RRID:AB_300798). To label PNNs, a solution of biotin-conjugated lectin Wisteria floribunda (WFA) (10 µg/ml; Sigma-Aldrich, L1516) was added in the primary antibody solution.

The secondary antibodies used in this study and their working concentrations are as follows: goat anti-chicken Alexa488 conjugated (1:1000; Abcam, ab150169), goat anti-rabbit Alexa633 conjugated (1:400; Life technologies, A21072), goat anti-mouse Alexa555 conjugated (1:1000; Cell Signaling, 4409S), goat anti-mouse Alexa647 conjugated (1:1000; Cell signaling, 4410S), goat anti-rabbit Alexa555 conjugated (1:400; Life technologies, A21430), goat anti-guinea pig Alexa647 conjugated (1:400; Life technologies, AB21450) and Alexa 568-conjugated streptavidin (1:500; Invitrogen, S-11226). All immunohistological experiments were performed on at least three different sections per brain region per animal.

### Confocal imaging

All images were acquired using Leica confocal microscopes (SP8 or SP8-STED). We imaged somatosensory (SSCX) and prefrontal cortex (PFC) layer 5 and basolateral amygdala (BLA) using 20X multi-immersion (NA 0.75) and 63X oil (NA1.4) objectives. We focussed on layer 5 of SSCX and PFC because this is where projection output neurons are located. In the PFC, we imaged the prelimbic region, because it plays a major role in regulating freezing behavior. The 20X objective was used to acquire images to analyze the percentage of: GFP^+^PV^+^/PV^+^ cells (recombination rate), GFP^+^PV^+^/GFP^+^ cells (Cre-recombination specificity), PNN^+^PV^+^/PV^+^ cells and Agg^+^PV^+^/PV^+^ cells. The 63X objective was used to acquire images to analyse the perisomatic innervation (number of perisomatic PV^+^ boutons and gephyrin^+^ punctas). At least three confocal images from three different brain sections were acquired per brain region with z-step size of 2 µm (20X) and 0.5 µm (63X). All the confocal parameters were maintained constant throughout the acquisition of an experiment.

### Image analysis

The number of positive cells (GFP^+^, PV^+^, PNN^+^ and/or Agg^+^) were manually identified and counted using ImageJ-Fiji software. To quantify the number of PV^+^, gephyrin^+^ and PV^+^/gephyrin^+^ punctas, images were exported as TIFF files and analyzed using Neurolucida software. PV^+^ boutons and gephyrin^+^ punctas were independently identified around the perimeter of a targeted cell after selecting the focal plane with the highest soma circumference. At least 4 perisomatic innervated somata were selected in each confocal image. Investigators were blind to the genotypes or conditions during the analysis.

### Behavioral testing

For all experiments, a camera was mounted above the arena; images were captured and transmitted to a computer running the Smart software (Panlab, Harvard Apparatus) or FreezeFrame software IMAQ 3 (Version 3.0.1.0). The sequence of animals tested was randomized by the genotype. Care was taken to test litters containing both the genotypes specific to the breeding.

#### Open Field

Each subject (P60) was gently placed at the center of the open-field arena and recorded by a video camera for 10 min. The recorded video file was analyzed with the SMART video tracking system (v3.0, Harvard Apparatus). The open field arena was cleaned with 70% ethanol and wiped with paper towels between each trial. The time spent in the center (45% of the surface) versus the periphery was calculated. Locomotor activity was indexed as the total distance travelled (m).

#### Elevated plus maze

The apparatus consisted of two open arms without walls across from each other and perpendicular to two closed arms with walls joining at a central platform. Each subject (P60) was placed at the junction of the two open and closed arms and allowed to explore the maze during 5min while video recorded. The recorded video file was analyzed with the SMART video tracking system (v3.0, Harvard Apparatus) to evaluate the percentage of time spent in the open arms (open/(open+closed) x 100) and the number of entries in the open arms as a measure of anxiety-related behavior.

#### Fear conditioning and fear extinction

Fear conditioning and extinction were evaluated in two different contexts (contexts A and B, respectively). The context A consisted of white walls, a grid floor, and was washed with 70% ethanol before and after each session. The context B consisted of black and white stripped walls, a white plexiglass floor, and was washed with 1% acetic acid before and after each session. Two different fear conditioning protocols have been used in this study, as previously published ^28^.

*PV-Cre*^*+/−*^*;Hdac2*^*lox/lox*^ males and their *PV-Cre*^*−/−*^*;Hdac2*^*lox/lox*^ or *PV-Cre*^*−/−*^*;Hdac2*^*lox/+*^ control males littermates, or *Hdac2*^*lox*^ wildtype mice, were conditioned using the following protocol. On day 1, mice have been allowed to freely explore the context A for habituation. On day 2, mice were conditioned using 5 pairings of the CS (CS duration 5 s, white noise, 80 dB) with a co-terminating US (1s foot-shock, 0.6mA, inter-trial interval: 30 – 60 s). On days 3 and 4, conditioned mice were submitted to extinction training in context B, which they were presented 12 CS at 30 s intervals on each day. Retrieval of extinction and context-dependent fear renewal were tested 7 days later in context B and A, respectively, using 4 presentations of the CS.

*PV-Cre*^*+/−*^*;Hdac2*^*lox/lox*^ males and their *PV-Cre*^*+/−*^*;Hdac2*^*+/+*^ control males littermates were conditioned using the following protocol. On day 1, mice have been allowed to freely explore the context A for habituation. On day 2, mice were conditioned using 5 pairings of the CS (CS duration 30 s, 50ms pips repeated at 0.9Hz, 2ms rise and fall, pip frequency: 7.5 kHz, 80 dB) with a US (1s foot-shock, 0.6mA, inter-trial interval: 30 – 150 s). The onset of the US coincided with the offset of the CS. On days 3 and 4, conditioned mice were submitted to extinction training in context B which they were presented 12 CS at 30 s intervals on each day. Retrieval of extinction and context-dependent fear renewal were tested 7 days later in context B and A, respectively, using 4 presentations of the CS.

Mice were excluded from analysis if they did not show effective fear extinction (EXT2 CS11-12 freezing more than 80% of time): Figure 1b: n=3 control mice excluded, n=0 *PV-Cre*^*+/−*^*;Hdac2*^*lox/lox*^ mouse excluded, Figure 1g: n=0 vehicle-injected *Hdac2*^*lox*^ mouse excluded, n=1 BRD6688-injected *Hdac2*^*lox*^ mouse excluded, Supplemental Figure 2: n=0 *PV-Cre*^*+/−*^*;Hdac2*^*+/+*^ mouse excluded, n=1 *PV-Cre*^*+/−*^*;Hdac2*^*lox/lox*^ mouse excluded. Mice were also excluded from analysis if they did not show effective fear acquisition (EXT1 CS1-2 freezing less than 15% of time): Supplemental Figure 2: n=0 *PV-Cre*^*+/−*^*;Hdac2*^*+/+*^ mouse excluded, n=1 *PV-Cre*^*+/−*^*;Hdac2*^*lox/lox*^ mouse excluded.

### Viral vector and stereotaxic injections

pAAV.EF1α-DIO-Flag-Hdac2_T2A_GFP (titre – 1E13 GC/mL) was cloned from pCIG-HDAC2^WT^-IRES-eGFP ^61^ and produced as AAV2/9 serotype by the Canadian Neurophotonics Platform, similar to the pAAV_EF1α_DIO_eYFP control virus (titre – 5E13 GC/mL [injected at 1:6 dilution]). *PV-Cre;Hdac2*^*+/+*^ or *PV-Cre;Hdac2*^*lox/lox*^ mice were used at PND 42-50 for all surgeries. Unilateral viral injections were performed at the following stereotaxic coordinates: +1.9mm Anterior-Posterior (from Bregma), +0.4mm Medio-Lateral and 1.6mm Depth from the cortical surface. Surgical procedures were standardized to minimize the variability of AAV injections. Glass pipette with the virus was lowered at the right depth and kept for 2 min before starting the injections. Each injection volume was kept to 46nl, and a total of 8 injections per mouse were done for each virus. To ensure minimal leakage into surrounding brain areas and to optimise viral spread in the desired area, injection pipettes were kept in the same position following the injection for 5 min. They were withdrawn at the rate of 0.25mm/min following the injection. Mice were kept for 3 weeks following viral injections to allow for optimal gene expression.

### Neuronal cell dissection and dissociation

Male mice aged P40-P43, were anesthetized with ketamine/xylazine (4:1), transcardically perfused with ice-cold aCSF solution of the following composition (in mM): 62.5 NaCl, 70 sucrose, 2.5 KCl, 25 NaHCO_3_, 1.25 NaH_2_PO_4_, 7 MgCl_2_, 1 CaCl_2_, 20 HEPES and 20 glucose. Brains were then quickly removed, transferred to ice-cold aCSF solution and fixed to a stage allowing to cut 300 μm slices with a VT1200 S, Leica vibratome. Slices were individually transferred to a cutting surface in ice-cold aCSF, where frontal cortex was micro dissected from the slices. Dissected tissue was then dissociated into cell suspension^62^ using the Papain dissociation system (Worthington) as per manufacturer’s instructions, with the addition of trehalose as described in ^63^. Post trituration, cells were then counted using iNCYTO C-Chip hemocytometers, and resuspended at 150-200 cells/μl in DropSeq Buffer (in mM): x1 HBSS, 7 MgCl_2_, 1 CaCl_2_, 20 glucose, 5% Trehalose and 10% BSA.

### Single cell capture and sequencing

Resuspended cells were run through a custom Drop-Seq setup allowing the capture of single cells and barcoded beads (ChemGenes) within nanoliter droplets as described in the original Drop-seq protocol (http://mccarrolllab.com/dropseq/)^64^. The setup closely resembled the original one^64^, with minor modifications. In particular, Medfusion 3500 syringe pumps (Smiths Medical) instead of KD Scientific Legato 100 pumps were used, as well as a different RNase inhibitor (SUPERase• In; catalog no. AM2694; Thermo Fisher Scientific).

Single-cell suspensions (150-200 cells/µl) were run alongside barcoded beads (140 beads µl/1) in 1.7 ml of cell lysis buffer (Tris 200mM, EDTA 20 mM, 20 % Sarkosyl, 20% Ficoll, DTT 50 mM; also containing 50 µl of RNase inhibitor), to produce beads attached to single-cell transcriptomes. Beads attached to single-cell transcriptomes were subsequently washed, reverse-transcribed, treated with exonuclease and counted using a Fuchs-Rosenthal hemocytometer (INCYTO C-Chip Cat-Nr:82030–474), as described in the original Drop-seq protocol^64^. Bead counting was performed using water and bead counting solution (10% polyethylene glycol, 2.5 M NaCl) in a 1:1 ratio as described in Gierahn et al. 2017^65^. For PCR amplification 4,000 beads (approximately 200 single-cell transcriptomes attached to microparticles) were used as input for each PCR. Individual PCR reactions were then pooled to achieve the desired number of single-cell transcriptomes attached to microparticles (STAMPS). In this experiment we used 2000 STAMPS which were cleaned up with AMPure XP (Beckmann Coulter Life Sciences) beads at a ratio of 0.6X prior to sequencing.

Correct size distribution and concentration of complementary DNA was determined using a Bioanalyzer High Sensitivity DNA assay (catalog no. 5067–4626; Agilent Technologies). Library preparation was performed using the Nextera XT DNA Library Preparation Kit (Illumina) with 600 pg input according to the Drop-seq protocol using Illumina i7 primers (N701—N706) together with a custom P5 primer (GCCTGTCCGCGGAAGCAGTGGTATCAACGCAGAGTAC) in the library PCR amplification (see the Drop-seq protocol^64^). Libraries were quality-controlled for size and concentration using the Bioanalyzer High Sensitivity DNA assay. Libraries were quantified using the Universal Library Quantification Kit (KAPA) before sequencing on a NextSeq 550 system (Illumina) at the Institute for Research in Immunology and Cancer (IRIC) in Montreal. Sequencing was performed on a NextSeq 500 system with the read settings R1 = 20 bp, R2 = 63 bp and index = 8 bp.

### Computational analysis of single-cell transcriptomic data

Demultiplexing of raw Illumina sequencing data was performed with bcl2fastq v.2.17 by the Institute for Research in Immunology and Cancer (IRIC) in Montreal. Fastq files were quality controlled and processed using the Drop-seq computational pipeline, similar to earlier report ^66^ (Drop-seq alignment cookbook v.1.2, January 2016) using Drop-seq tools v.1.12, to produce digital gene expression (DGE) matrices for each sample^64^. DGE matrices were processed in R (3.5.2) using Seurat (v.2.2.0) ^67,68^. Cells with less than 250 genes expressed and genes detected in less than 3 cells were filtered out from further analysis. Cell cycle scores for filtered cells were calculated using the function CellCycleScoring from Seurat with cell cycle gene lists from Nestorowa et al.^69^(https://satijalab.org/seurat/articles/cell_cycle_vignette.html). Expression values were normalized and scaled to 10000 transcripts with Seurat’s Normalize function. Normalized expression values were scaled using the ScaleData function from Seurat, regressing out the number of UMIs, Mouse ID, percent mitochondrial genes as well as S and G2M scores from cell cycle prediction. Wild-type cells from different experiments were further aligned with using canonical correlation analysis (CCA)^68^ using the RunMultiCCA using the top 1000 overlapping highly variable genes between samples function followed by AlignSubspace function in Seurat, using the experiment as a grouping variable with 15 cca dimensions. Dimensions selection was decided after visual inspection of the inflection point of the elbow plot generated by Seurat ElbowPlot() function. Dimensional reduction was performed using RunTSNE with 15 cca dimensions to represent cells in 2 tSNE dimensions. Clusters were identified using the FindClusters function on the first 15 aligned cca dimensions with a resolution parameter of 1.2. Cells were subsequently plotted in 2 tSNE dimensions to and biological cell types annotated manually based on known markers for brain cell types. Violin Plots were plotted using the VlnPlot function from Seurat with normalized gene expression values.

### Cell type Identification

Final cell types were attributed based on the identified top markers in each cluster. Clusters containing VGLUT1 were considered to be pyramidal Neurons, while clusters containing GAD1 and GAD2 were considered to be GABAergic interneurons (Figure 3a, Supplementary Figure 4). For more in depth analysis of neuron types as well as identification of non-neuronal cell subtypes, we took the top 30 and 50 markers from each cluster and compared them to the DropVis database^70^. The number of markers shared for each cluster was then calculated, and a percentage identity with DropViz clusters calculated.

### Fluorescent multiplex RNAscope

To evaluate *Acan* mRNA expression in naive condition, mice of both sexes were cervically dislocated, brains were dissected, fast-frozen in cold isopentane (~-70°C) and conserved at −80°C. To evaluate *Acan* mRNA expression after extinction training, adult males were cervically dislocated 3 hours after the end of extinction training on day 4 (late extinction).

Coronal brain sections (20 µm) were cut using a cryostat (Leica Microsystems) and mounted on Superfrost Plus Gold Glass Slides (Fisher Scientific, 22-035-813). Slides were subsequently stored at −80°C. Probes against *Acan* mRNA (439101), which codes for aggrecan, and *Pvalb* mRNA (421931-C2), which codes for PV, as well as other reagents for ISH and DAPI labelling, were purchased from Advanced Cell Diagnostics (ACD). Tissue pre-treatment, hybridization, amplification and detection were performed according to the *RNAscope Sample Preparation and Pretreatment Guide for Fresh Frozen Tissue using RNAscope Fluorescent Multiplex Assay* manual (ACD). During RNAscope hybridization, positive probes (320881), negative probes (320871), and *acan*/*Pvalb* probes were processed simultaneously. Briefly, the slides were removed from −80°C and immediately post-fixed in 4% PFA/PB for 15 min before dehydration in 5 min consecutive baths (70% EtOH, 50% EtOH, 2 × 100% EtOH). The slides were air dried and hydrophobic barrier was created around each section with ImmEdge Pen (310018). Protease IV (322336) was added to each section and incubated at room temperature for 30 min followed by two washes in distilled water. For detection, probes were added to each section and incubated for 2h at 40°C. Unbound probes were subsequently washed away by rinsing slides in 1X wash buffer (310091) 2 × 2 min. AMP reagents (AMP1 (320852), AMP2 (320853), AMP3 (320854), AMP4A (C1 probes-Alexa-488 and C2 probes-Atto-550) (320855)) were added to each section and incubates for as per the manufacturer’s instructions (*RNAscope Fluorescent Multiplex Reagent Kit PART 2 Manual*), and washed in wash buffer for 2 × 2min. Sections were stained with DAPI (320858) for 30 s, and then mounted with Prolong Diamond Antifade Mountant (Invitrogen P36961). Images were acquired with a SP8-STED confocal microscope (Leica Microsystems), using a 63X (NA 1.4) objective, voxel size: 0.06µm x 0.06µm x 0.299µm, and deconvolved using Huygen’s express deconvolution option. These experiments were performed using tissue from three different mice for each genotype and/or treatment. To determine the number of *Acan* dots per PV^+^ cell, images were analysed using a custom-made ImageJ-Fiji macro. Briefly, a dark region was selected on each focal plane to measure the mean grey value (MGV) as background (BCKG). Then, for each channel and each focal plane, the BCKG value was subtracted, and intensity value of each pixel was readjusted by multiplying it with 65535/(65535-BCKG). Then, each PV^+^ cell was manually selected, filtered with the Gaussian Blur 3D option and analyzed using the 3D objects counter (threshold = 4500, minimal voxel size = 6). Results are presented as mean of the number of objects per PV^+^ cell ± s.e.m.

### Drug treatment

BRD6688 (Glixx Lab, #GLXC-05908) was dissolved in DMSO (1% of the total resultant solution) and then diluted in 30% Cremophor/69% in physiological saline (H_2_O containing 0.9% NaCl (Hospira, 04888)), for a final dosage solution of 1 mg/kg. Vehicle solutions consisted of the aforementioned solution without the compound. Solutions were prepared immediately before injection and administered to either *Hdac2*^*lox*^ wildtype or *PV-Cre*^*+/−*^*;Hdac2*^*lox/lox*^ via intraperitoneal injection 6 hours prior to extinction training.

### siRNA-delivering peptides

TARBP-BTP RNA-binding protein was produced by the National Research Council Canada (NRC), Montreal, Quebec, Canada, as described previously^42^. The original pET28a-His-pTARBP-BTP coding plasmid was provided by CSIR-CCMB. TARBP-BTP:siRNA complex were prepared as previously described^42,43^. Silencer Select *Acan* mouse targeting siRNA (Life technologies, #s61116) and Silencer Select Negative Control siRNA (Life technologies, #4390843) were resuspended at 200 µm with nuclease-free water. For in vivo delivery, TARBP-BTP and siRNA were drop wise mixed together at 5:1 mole ratio. The complex was incubated 20 min at room temperature and administered intravenously (i.v.) via the lateral tail vein at a dose of 20 nmol:4 nmol TARBP-BTP:siRNA in a total volume of 150 µL per mouse. 48h after injection, 3 P60 *Hdac2*^*lox*^ mice injected with siCtrlNeg and 4 P60 *Hdac2*^*lox*^ mice injected with siAcan were cervically dislocated, brains were dissected and fast-frozen in cold isopentane (#x007E;-70°C) and conserved at −80°C before proceeding with Fluorescent multiplex RNAscope. To investigate Acan knockdown effect on fear extinction, 21 Hdac2lox were fear-trained using the following protocol. On day 1, mice have been allowed to freely explore the context A for habituation. On day 2, mice were conditioned using 5 pairings of the CS (CS duration 5 s, white noise, 80 dB) with a co-terminating US (1s foot-shock, 0.6mA, inter-trial interval: 30 – 60 s). On day 3, 11 mice were i.v. injected with TARBP-BTP:siCtrlNeg and 10 mice were i.v. injected with TARBP-BTP:siAcan at a dose of 20nmol:4nmol per mouse. On days 4 and 5, conditioned mice were submitted to extinction training in context B, which they were presented 12 CS at 30 s intervals on each day. Retrieval of extinction and context-dependent fear renewal were tested 7 days later in context B and A, respectively, using 4 presentations of the CS.

### Statistical analysis

Data were systematically tested for normal distribution, with the D’Agostino & Pearson omnibus normality test. Differences between two groups of normally distributed data with homogenous variances were analyzed using parametric student’s t-test, while not normally distributed data were analyzed with the Mann-Whitney test. To evaluate genotype effects on PV^+^Gephyrin^+^ perisomatic puncta in naive mice and in mice following extinction training, 2-ways ANOVA test was performed followed by Sidak *posthoc* test. To evaluate the effect of the virally-expressed *Hdac2* in *PV-Cre*^*+/−*^*;Hdac2*^*lox/lox*^ on Aggrecan and PNN, Kruskall-Wallis tests were performed followed by Dunn’s multiple comparison test. For fear conditioning analysis, we evaluated the extinction rate by Repeated measures two-ways ANOVA, while freezing time at extinction retrieval and fear renewal tests were analyzed with unpaired two-tailed t-test, if normally distributed and Mann-Whitney if not. Results were considered significant for values of *P*<0.05. Data are presented as mean ± standard error of mean.

### Data availability

All data are available upon request.

## Acknowledgements

We thank Dr. Elsa Rossignol for her insightful suggestions and critical reading of the manuscript. We would like to thank Josianne Nunes-Carriço, Antônia Samia Fernandes do Nascimento and Antoine Farley for their technical assistance. We acknowledge the Comité Institutionnel de Bonne Pratiques Animales en Recherche (CIBPAR), and all the personnel of the animal facility of the Research Center of CHU Sainte-Justine (Université de Montreal), as well as Dr. Elke Küster-Schöck at the plateforme d’imagerie microscopique (PIM) for instrumental technical support. We thank J. Huber at the IRIC genomic platform for performing the Illumina sequencing for the Drop-seq libraries and P. Gendron at the IRIC Bioinformatics platform for data demultiplexing. We thank the Canadian Neurophotonic Platform for viral production. We also would like to thank Andrei Vlad Varlan, Kayleigh Gauvin and Théo Badra for their help with data analysis, CCMB’s central facility for plasmid DNA sequencing, and the National Research Council Canada for TARBP-BTP carrier protein production and purification.

## Fundings

This work was supported by the Canadian Institutes of Health Research (G.P., G.DC), Natural Sciences and Engineering Research Council of Canada, EraNet-Neuron (G.DC). M. L-J was supported by le Fonds de Recherche du Québec en Santé (FRQS).

### Author contributions

MLJ, GDC designed the experiments. MLJ, BC, PC, DC, SL, FD, DR performed the experiments. MLJ, PC, FW, FD, DR, HA, GP and GDC analysed data. KS, VG, ABP, GA and GP provided critical technical support. MLJ, GDC wrote the manuscript. All authors read and corrected the manuscript.

