## Supplemental Figures for "Enhancing adult neuroplasticity by epigenetic regulation of Parvalbumin-expressing GABAergic cells"

### Supplemental Material

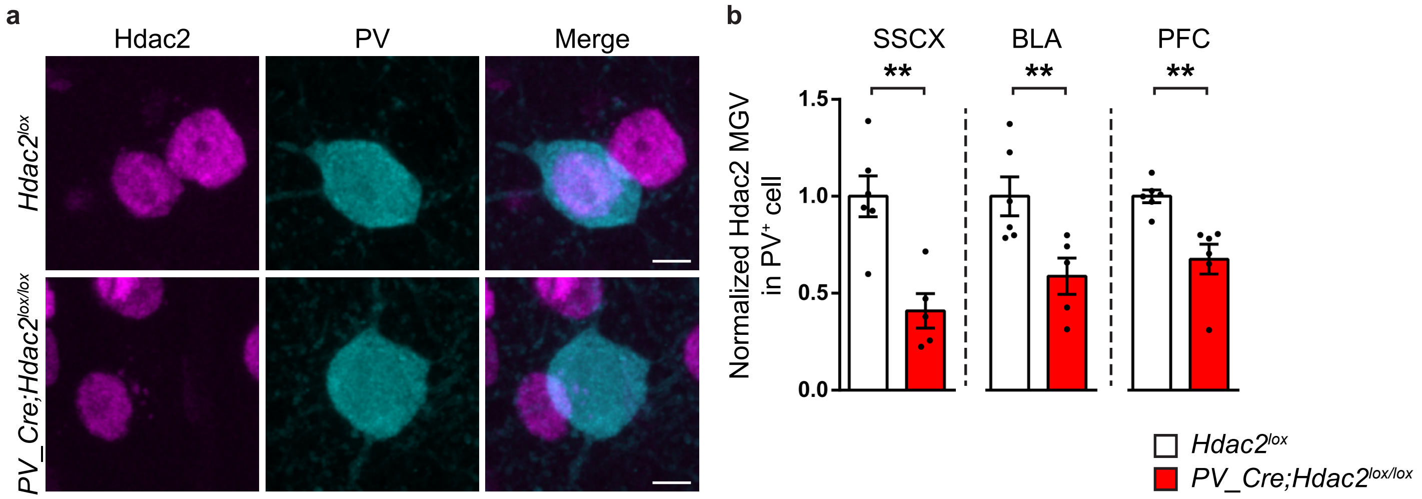


**Supplementary Figure 1. Hdac2 expression is decreased in PV+ cells of *PV_Cre;Hdac2lox/lox* mice.**

(**a**) PFC coronal sections from P60 *PV_Cre;Hdac2lox/lox* and control littermates immunolabeled for Hdac2 (magenta) and PV (cyan). Scale bar, 5 µm. (**b**) Mean normalized fluorescent intensity of Hdac2 signal in PV+ cell nuclei is reduced in conditional knockout mice compared to control littermates in somatosensory cortex (SSCX, Mann-Whitney test, *P*=0.0087), basolateral amygdala (BLA, Mann-Whitney test, *P*=0.0087) and prefrontal cortex (PFC, Mann-Whitney test, *P*=0.0022). Number of mice: PFC: *Hdac2lox/lox* n=6; *PV_Cre;Hdac2lox/lox* n= 6, BLA and SSCX: *Hdac2lox/lox* n=6; *PV_Cre;Hdac2lox/lox* n= 5. MGV = mean grey value. Graph bars represent mean ± s.e.m. Circles represent individual mouse values.  ** *P*<0.01.

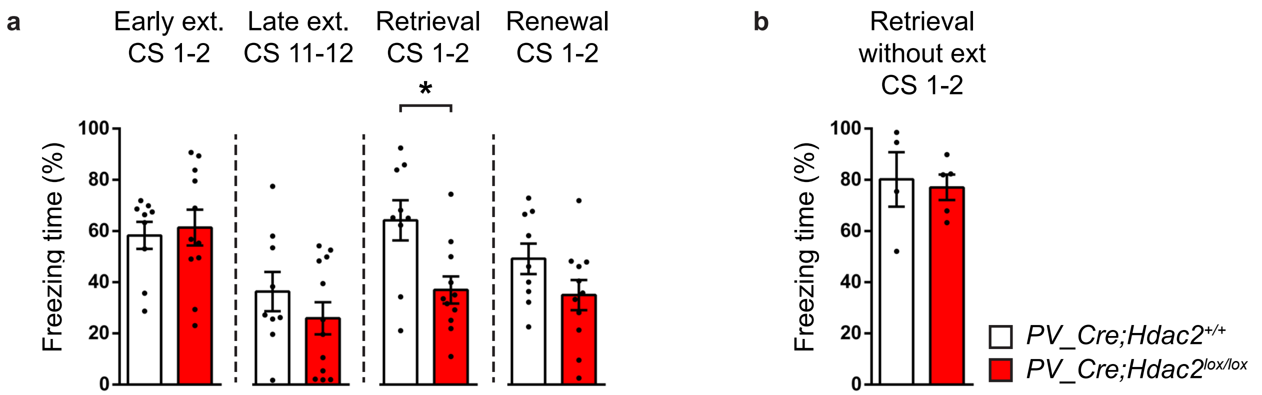


**Supplementary Figure 2. Adult *PV_Cre;Hdac2$^{lox/lox}$* mice show increased fear extinction retention compared to their control *PV_Cre;Hdac2$^{+/+}$* littermates.**

(**a**) Both *PV_Cre;Hdac2$^{+/+}$* and *PV_Cre;Hdac2$^{lox/lox}$* mice show efficient fear extinction. Repeated two-way ANOVA; $F_{genotype}$ (1,19)=0.1393, *P*=0.7131, $F_{extinction}$(11,209)=7.496, *P*<0.0001, $F_{genotype*extinction}$ (11,209)=1.079, *P*=0.3800. One week after extinction training, *PV_Cre;Hdac2$^{lox/lox}$* mice show significantly reduced freezing time than *PV_Cre;Hdac2$^{+/+}$* mice in the fear retrieval test (unpaired two-tailed t-test, *P*=0.0064), but not in the fear renewal test (unpaired two-tailed t-test, *P*=0.1080). Number of mice: *PV_Cre;Hdac2$^{+/+}$* n=9, *PV_Cre;Hdac2$^{lox/lox}$* n=12. (**b**) In the absence of extinction training, *PV_Cre;Hdac2$^{lox/lox}$* mice show similar freezing time as their control littermates 10 days after fear training (Mann-Whitney test, *P*=0.7143). Number of mice: *PV_Cre;Hdac2$^{+/+}$* n=4, *PV_Cre;Hdac2$^{lox/lox}$* n=5. Graph bars represent mean ± s.e.m. Circles represent individual mouse values. * *P*<0.05.

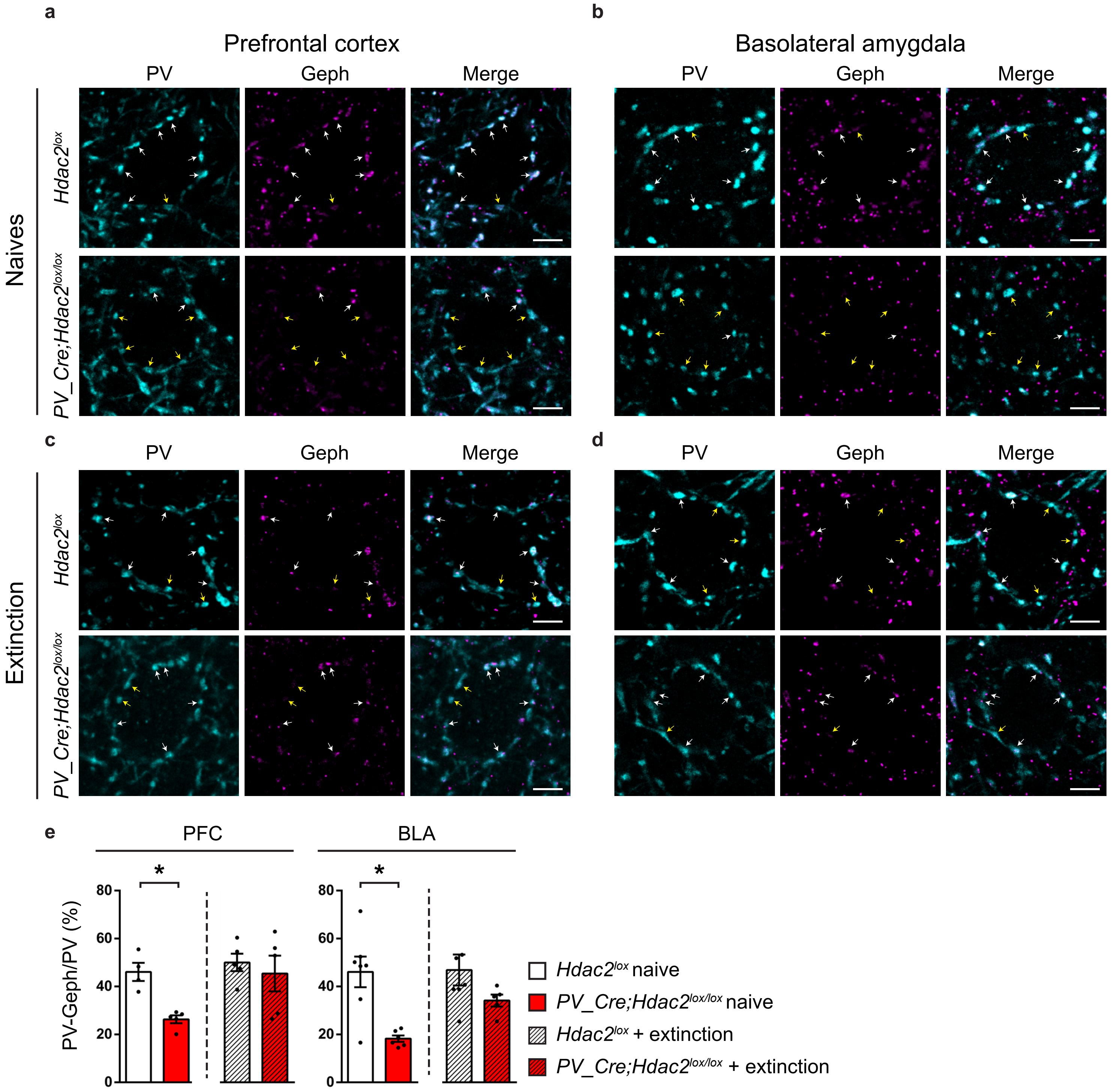
a
Prefrontal cortex
PV
Geph
Merge
b
Basolateral amygdala
PV
Geph
Merge
Naives
Hdac2lox
PV_Cre;Hdac2lox/lox
c
PV
Geph
Merge
d
PV
Geph
Merge
Extinction
Hdac2lox
PV_Cre;Hdac2lox/lox
e
PFC
BLA
*
*
PV-Geph/PV (%)
80
60
40
20
0
Hdac2lox naive
PV_Cre;Hdac2lox/lox naive
Hdac2lox + extinction
PV_Cre;Hdac2lox/lox + extinction

**Supplementary Figure 3. PV$^+$ cell perisomatic connectivity show increased remodeling following extinction training in PFC and BLA of *PV-Cre;Hdac2$^{lox/lox}$* mice.**

(**a-d**) Coronal brain sections of PFC (**a, c)** and BLA (**b, d)** immunolabeled for PV (cyan) and gephyrin (magenta) (**a, b**) Naive *PV-Cre;Hdac2$^{lox/lox}$* and control littermates. (**c, d**) *PV-Cre;Hdac2$^{lox/lox}$* and control littermates 24h after late extinction training. Scale bar, 10 μm. (**e**) The percentage of perisomatic PV$^+$ boutons co-localizing with gephyrin is significantly different between the two genotypes in naive condition but not 24 hrs after extinction training. PFC: two-way ANOVA; $F_{genotype}$ (1,15)=6.564, P=0.0217, $F_{extinction}$( 1,15)=5.861, P=0.0286, $F_{genotype*extinction}$(1,15)=2.523, P=0.1331. BLA: two-way ANOVA; $F_{genotype}$ (1,21)=14.56, P=0.0010, $F_{extinction}$ (1,21)=2.478, P=0.1304, $F_{genotype*extinction}$ (1,21)=2.018, P=0.1701. Sidak's posthoc test show statistical significance in PFC and BLA between *PV-Cre;Hdac2$^{lox/lox}$* and control littermates only in naive condition (PFC: naives: P=0.0241, extinction: P=0.7386; BLA: naives: P=0.0021, extinction: P=0.2140). Mice numbers: PFC: Naives control n=4, *PV-Cre;Hdac2$^{lox/lox}$* n=5; Extinction control n=5, *PV-Cre;Hdac2$^{lox/lox}$* n=5. BLA: Naives control n=7, *PV-Cre;Hdac2$^{lox/lox}$* n=6; Extinction control n=7, *PV-Cre;Hdac2$^{lox/lox}$* n=5.

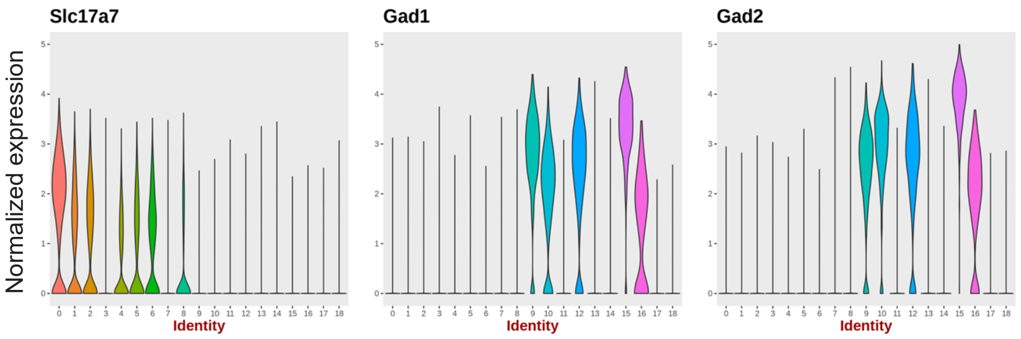


**Supplementary Figure 4. Expression levels of VGLUT1 (Slc17a7), GAD1 (Gad1) and GAD2 (Gad2) in clusters 1-18.** Expression levels are shown as normalized gene expression for each cluster. Cells containing VGLUT1 are considered as glutamatergic neurons, cells containing GAD1 and GAD2 are described as GABAergic neurons. Cells without these markers were labeled as non-neuronal.

**Related to Figure 3.**

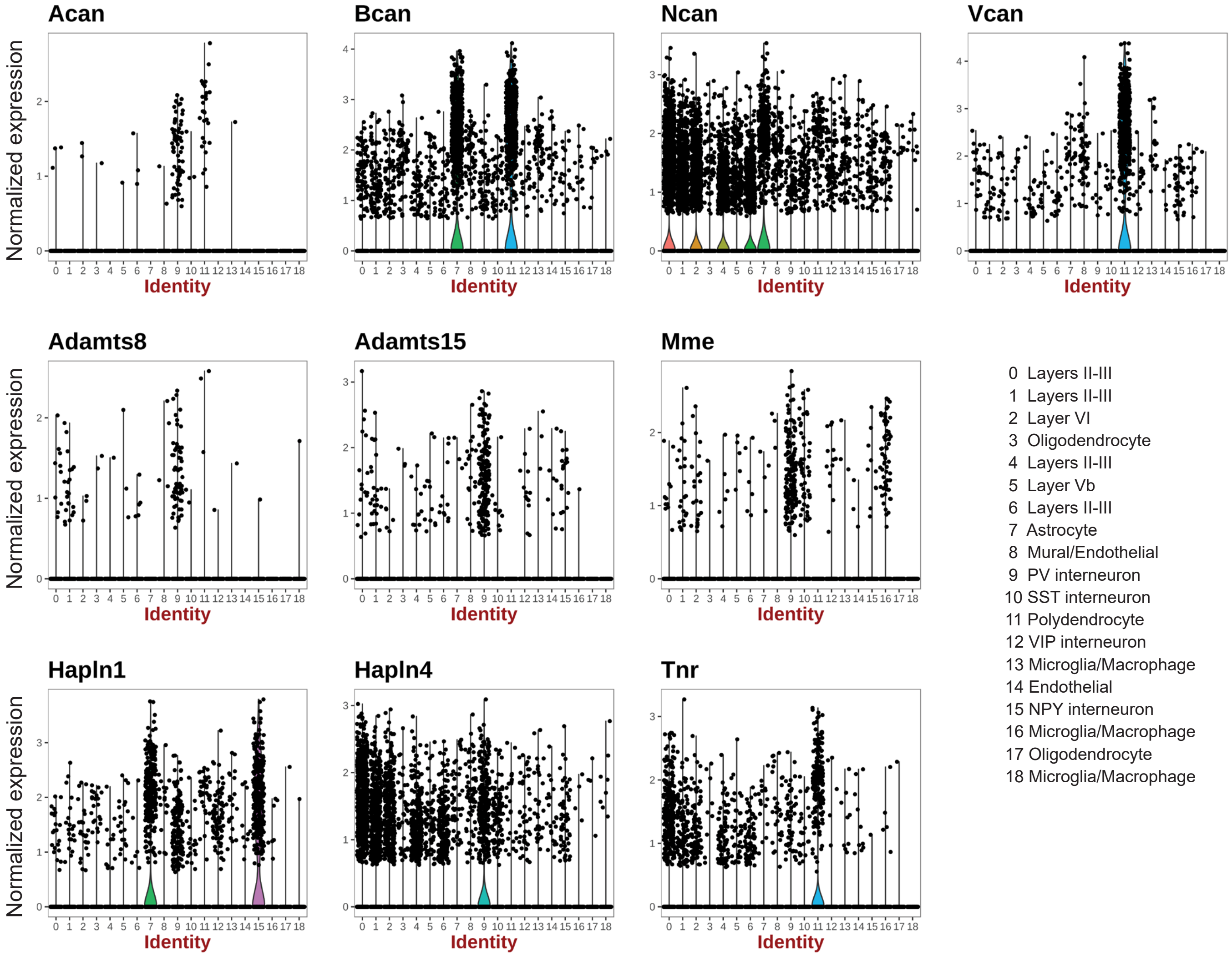


**Supplementary Figure 5. mRNA expression of different PNN components and metallo-proteases in distinct cell types in prefrontal cortex of P40 mice identified by single cell RNAseq (DropSeq).** Data is shown as violin plots, where clusters are graphed against normalized expression levels for each gene.

**Related to Figure 3.**
*Acan* expression is mostly restricted to $PV^{+}$ cells and, to a lesser extent, polydendrocytes.

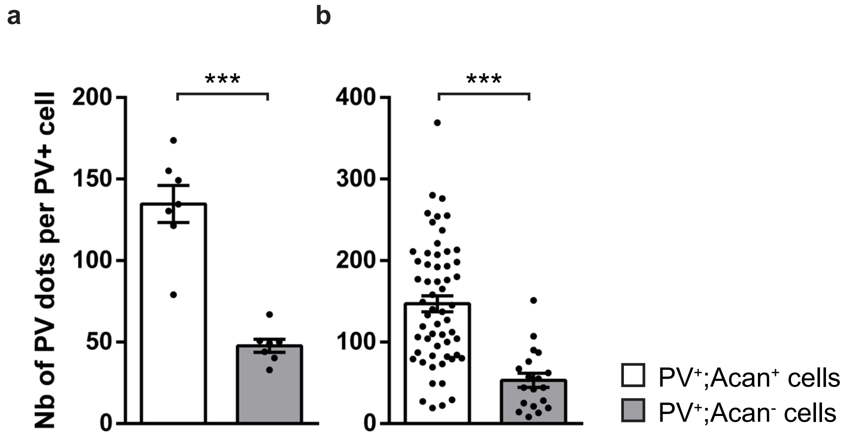


**Supplementary Figure 6. Prefrontal cortical PV$^+$ cells with detectable levels of *Acan* mRNA express more *Pvalb* mRNA dots than PV$^+$Acan$^-$ cells.** (a) Mean of *Pvalb* mRNA dot numbers in PV$^+$Acan$^+$ and PV$^+$Acan$^-$ cell somata averaged per animal. Unpaired t-test with Welch's correction, p=0.0001. N=7 control mice. (b) Number of *Pvalb* mRNA dots in PV$^+$Acan$^+$ and PV$^+$Acan$^-$ cell somata for each analysed cell. Unpaired t-test with Welch's correction, p<0.0001. N=58 PV$^+$Acan$^+$ cells, N= 19 PV$^+$Acan$^-$ cells. Graph bars represent mean ± s.e.m. Circles represent individual values.*** p<0.001
